# Renovating the Barnes maze for mouse models of Dementia with STARR FIELD: A 4-day protocol for learning rate, retention, and cognitive flexibility

**DOI:** 10.1101/2024.11.30.625516

**Authors:** Aimee Bertolli, Oday Halhouli, Matthew A. Weber, Yiming Liu-Martínez, Linder H. Wendt, Brianna Blaine, Ramasamy Thangavel, Sura Smadi, Preston J. Pellatz, Danlin Liu, Kaisa N. Bornhoft, Kalpana Gupta, Riley Dean, Margaret Tish, Gemma Kerr, Shane A. Heiney, Eric B. Emmons, Qiang Zhang, Serena B Gumusoglu, Nandakumar S. Narayanan, Joel C. Geerling, Georgina M Aldridge

**Author notes:** Corresponding Author Georgina M Aldridge, 169 Newton Road, Pappajohn Biomedical Discovery Building-5334 University of Iowa, Iowa City, 52242. **Conflict of Interest:** There are no conflicts of interest.

## Abstract

Land-based spatial mazes are a low-stress method to evaluate learning in rodent models of dementia. By using innate exploratory and hiding behavior, the Barnes maze requires fewer trials, allowing examination of early learning rate and retention, as well as executive and motivational features that can be characteristic of non-amnestic dementias. However, unwanted odor cues may disrupt interpretation of acquisition rate during typical learning trials. We designed and tested our Barnes FIELD protocol (Find the Invisible Exit to Locate the Domicile) to improve reproducibility, allow evaluation of learning trials, and limit experimenter influence. The protocol uses 3D-printed escape shuttles and docking tunnels, allowing mice to exit the maze to the home cage. We show evidence that our shuttles mitigate the possibility of undesired cues. We demonstrate the feasibility of our protocol across several models of cognitive impairment and aging, and develop an additional stage, the STARR (Spatial Training And Rapid Reversal) maze, to better challenge behavioral flexibility. By examining commonly used outcome measures we identify important considerations for interpretation. These insights are used to evaluate several models of cognitive change, including deficits in an Alzheimers disease mouse model, and behavioral flexibility in a model of brainstem dysfunction. This work provides comprehensive instructions to build, perform, and analyze a robust spatial maze that expands the range of behavioral and motivational outcomes that can be identified and screened. Our findings will aid interpretation of traditional protocols, enhance rigor and reproducibility, and provide an updated method to screen for cognitive changes in mice.

## INTRODUCTION

Neurodegenerative dementias are devastating incurable diseases that affect 55 million people worldwide and cause severe disability and death [1]. Mouse models are used for studying mechanisms and testing new therapies; they have similar cortical architecture, many conserved brainstem projections, and abundant available genetic tools [2–5].

Behavioral tasks in rodents are widely used to evaluate short-term memory deficits in models of Alzheimer’s dementia (AD). However, fewer studies focus on non-memory aspects of cognition, such as executive dysfunction, that are common features of other forms of dementia, including frontal variant AD, Lewy body dementias, frontotemporal dementias, vascular dementia, and normal pressure hydrocephalus [6–9].

Assays that use innate motivation like novelty and memory of an escape location (e.g. novel object recognition, Morris Water Maze) do not require extensive training [10–12]. In these innate tasks, short-term memory retention is often used as the key outcome measure [13, 14]. However, naturalistic tasks also offer the opportunity to test active acquisition and motivation, rather than only retention. This includes outcomes impacted by executive function (e.g., learning rate, working memory, behavioral flexibility, sequencing, planning) and motivation (e.g., perseveration, perseverance, exploration, anxiety). Both motivation and executive dysfunction can be key features of neurodegenerative diseases [15, 16].

Studies probing executive functions often require extended training protocols, expensive operant chambers, or strong motivators [17]. To improve our ability to detect a wider range of behavioral phenotypes, we chose to optimize the Barnes Maze, a widely used spatial learning and memory task originally developed by Carol Barnes for rats[18]. This land-based maze is used in hundreds of published research articles yearly. Specific benefits are that it: 1) resembles naturalistic rodent behavior (exploration and remembering hiding locations); 2) does not require food or water motivation; 3) is comparatively low-stress; 4) can be adapted for mice with motor deficits (no forced swimming), and 5) is widely available and relatively inexpensive to build [17, 19–21].

In this task, mice are placed atop the center of a large, bright, open platform with holes around the outer edge (Fig. 1). One of the holes leads to a dark escape box. Mice have an innate desire to find escape routes and remember their location [12]. Investigators use various outcome measures, such as distance, latency, or number of explored holes during learning trials as a measure of learning rate. Secondly, a separate probe trial is performed. During the probe, the escape box is removed. An increased proportion of time exploring the location where the target was located is used as evidence of memory for that location.

**Figure 1:**
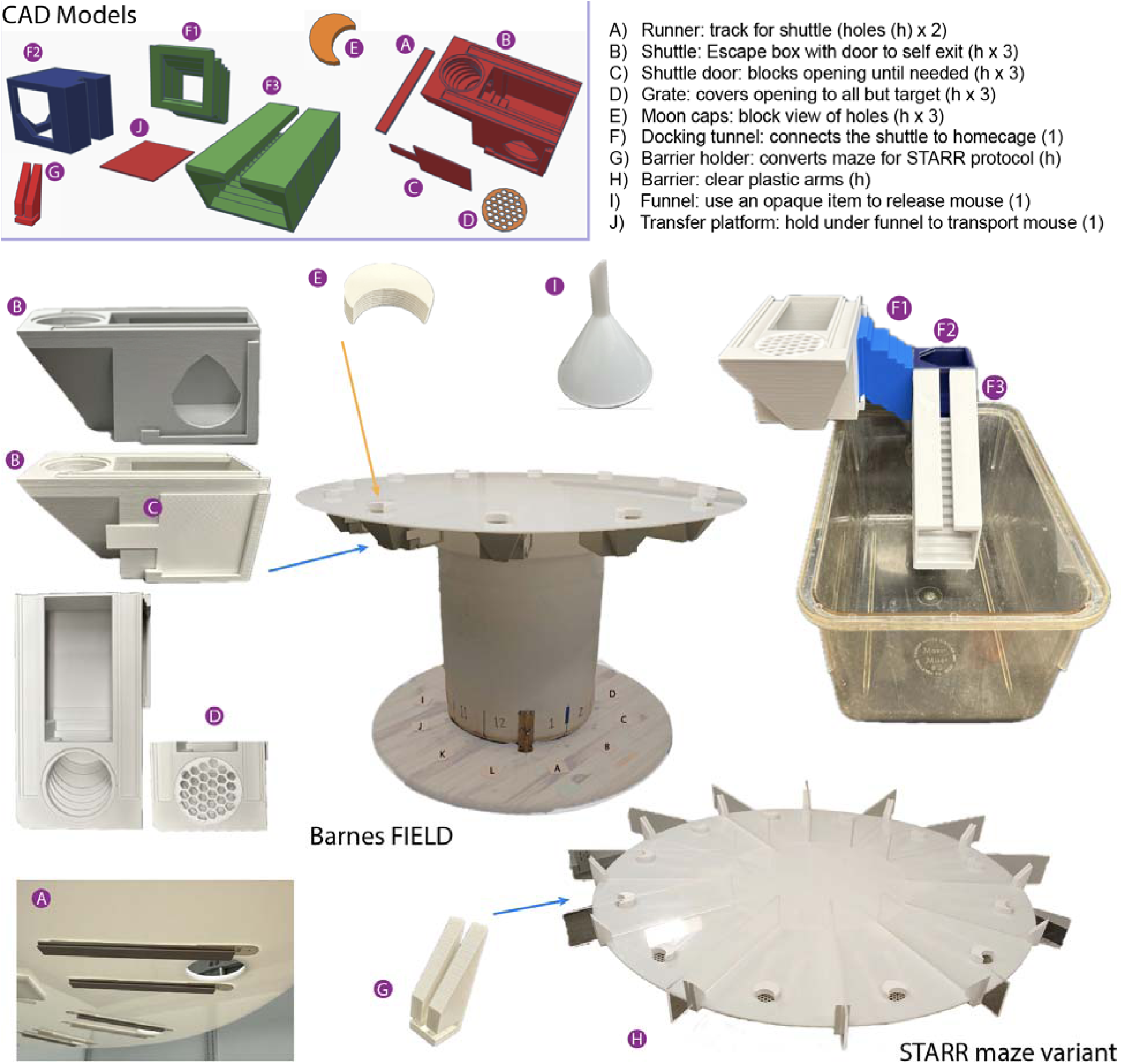
Barnes FIELD equipment: All 3D printed items can be printed without supports. Minor assembly is required only for docking tunnel (F1-F3). Shuttles (B), with or without blocking grates (D) slide onto runners (A) that are positioned under a traditional spatial platform. All holes should have a clean, dry shuttle, with only the target shuttle open. Moon caps (E) can be positioned to block visual cues from the center of the maze. The mouse is released onto the maze using a funnel (I) with a thin rigid plastic sheet to transfer the mouse (J). Only AFTER the mouse enters the shuttle, the docking tunnel is attached (F), the home cage is positioned underneath, and the door between (C) is opened to allow the mouse to self-exit. If the mouse remains in the tunnel after its reward period, the tunnel contains holes to help with gently guiding it out. **STARR maze equipment:** Plastic, transparent barriers (H) are added to create a radial style maze using printed barrier holders (G). The minimum recommended copies to run the task are listed, additional copies are helpful in case of breakage.

During our early use of the Barnes maze, we noted that unintended olfactory cues can confound learning trials and inadvertently impair interpretation. We noted that if the mouse’s home-cage was attached to the escape box, some mice could repeatedly find the target hole without approaching any other hole, even when the target was moved at random every trial. Most commercial devices use a single escape box for all learning trials, which is also a potential source of olfactory cues. Although the escape box is cleaned after each trial in most Barnes protocols, its solitary nature represents a clear olfactory confound. This confound leads many researchers to analyze only the probe trial, for which the escape box is removed.

Limiting analysis to the probe trials misses evaluation of learning trials necessary to evaluate learning rate, search strategy, exploration efficiency, and response to target reversal for evaluation of behavioral flexibility. Therefore, we designed an inexpensive, 3D-printable escape box (“shuttle”) that could be deployed on most standard Barnes mazes, allowing researchers to have multiple copies of the escape box to mitigate the confound of intramaze olfactory cues. The adapted protocol, Barnes FIELD (Find the Invisible Exit to Locate the Domicile), creates exits that are “invisible,” and shuttles designed to link to the home cage (after the exit is located), providing naturalistic reinforcement and reduced experimenter interference.

We directly tested the benefit of the Barnes FIELD protocol by comparing unwashed shuttles to clean, 3D-printed shuttles. We demonstrate mice were able to detect an olfactory cue based on probe trial outcomes. The design of the study also allowed us to test recent evidence showing that most noticeable gains in latency and distance during early learning trials are driven by non-spatial learning [22–24]. In this context, non-spatial learning may include improvements secondary to meta-learning, motivation, procedural-learning, and habituation that are acquired simultaneously with spatial location. Therefore, we further adapted the spatial Barnes maze to incorporate intermittent probe trials across learning to isolate spatial memory acquisition and retention. Finally, we developed an additional stage, the STARR maze protocol (Spatial Training And Rapid Reversal), that can be incorporated at the end of any Barnes protocol (including our Barnes FIELD protocol) to screen for behavioral flexibility.

These findings allowed us to test our Barnes FIELD and STARR maze protocols in models of aging and dementia currently used in neurodegenerative laboratories at the University of Iowa. These include: 1) a model of beta-amyloid plaque formation, 5xFAD [25];, 2) a model of normal-pressure hydrocephalus induced by cisternal injection of kaolin [26];, 3) a model of differential aging comparing virgin and postpartum one-year old female mice [27]; and 4) a model of locus coeruleus damage (DSP-4 toxin), mimicking the norepinephrine dysfunction seen in Lewy body disease.

We evaluate the use of both the Barnes FIELD and STARR maze protocols in these models, highlighting aspects of the task that can be evaluated when all trials can be more confidently interpreted.

## MATERIALS AND METHODS

### Equipment

For this study, all items were 3D printed in Polylactic acid (PLA), without supports, using a Prusa i3 MK3S+ (Fig. 1). Grey PLA (AmazonBasics, Color: Light Grey, 1.75 mm) was used for shuttle boxes based on early testing; as white allowed too much light into the shuttle boxes, and darker colors interfered with tracking software. White PLA (AmazonBasics, Color: White, 1.75 mm) was used for moon caps, doors, grates and barrier holders (Fig. 1). STL files are available at TinkerCad (Autodesk) and at aldridge.lab.uiowa.edu/open-science. The original PLA used for printing is no longer available, but any PLA or PLA+ of light grey and white, respectively, should be equivalent. However, identical items should be printed with the same material; we recommend printing extras. <u>Shuttle boxes, moon caps, grates, and runners</u> do not require assembly.

<u>Docking tunnels</u> (Fig. 1F) were printed in three pieces and glued with Loctite® Instant Mix Epoxy (47 oz; Henkel Corp., U.S.A.).). <u>Runners</u> were attached under the platform using command strips (Command™ Large Picture Hanging Strips; 3M, U.S.A.). To assemble, place the runners into the shuttles then add command strips along the top of each runner. Allow to set per manufacturer guidelines, then remove backing from other side of strip. Align the shuttle on the underside of the maze, aligning the opening of the shuttle with the 2-inch platform hole, and press firmly. The shuttle should slide easily off and onto the maze like a drawer, allowing for easy cleaning and replacement.

### Platform and rotating maze

A commercial Barnes maze platform set can cost thousands of dollars and usually includes only a single escape box. However, many labs already own this apparatus or parts of one that can be adapted with the Barnes FIELD shuttles. The original maze available to our laboratory (brand unknown) had only the white plastic platform: 122 cm (∼48 inches) in diameter, 0.635 cm (1/4 inch) thick, with 40 holes (2 inches/5.08 cm in diameter), equally spaced around the perimeter. It was missing all other parts. All but 10 of the 40 holes were covered by plastic disks from the underside until a dedicated platform could be built. For the 12-hole maze (DSP-4), the platform was custom cut by a local plastic manufacturer.

We recommend investigators that do not have their own maze to custom order the top out of high density polyethylene (HDPE) from any plastic manufacturer (Supplemental Figs. 1 and 2). Based on our early internal testing of 8-vs. 10-vs. 12-vs. 20-hole mazes, and published data in the literature [28] we did not find improved ability to discriminate between groups of mice with more holes (data not shown). Past a certain point (e.g., 32 holes), mice may find the maze more difficult, requiring more learning trials and risk of mice switching to serial search strategies [29]. We recommend making a maze with 12 holes, named by the hands of a clock (Supplemental Fig. 2), which can be divided into classic quadrants with one target hole and two adjacent holes.

**Figure 2:**
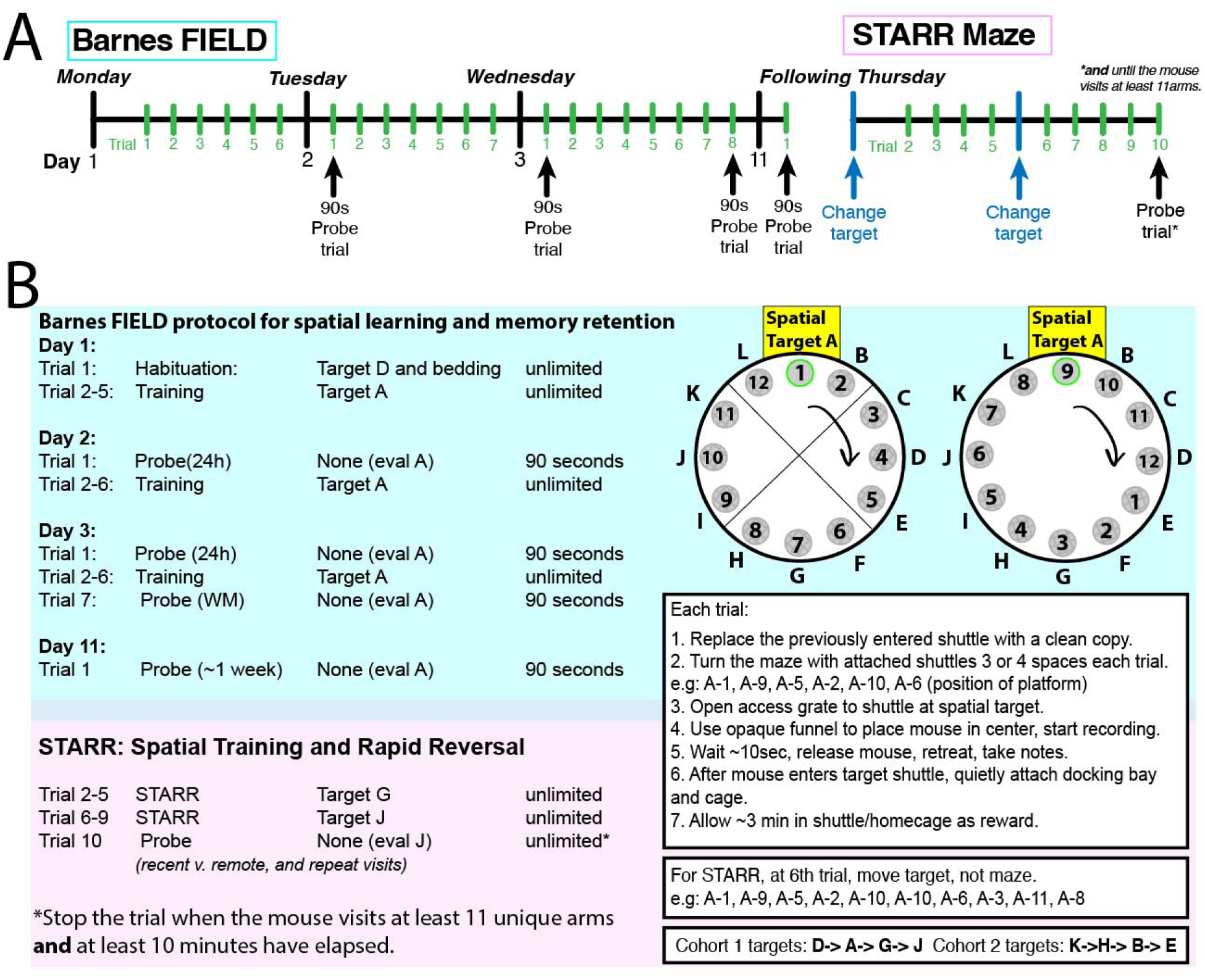
Printable Short Form Protocol. (A) A 4-day protocol for spatial acquisition, short-and long-term memory (Barnes FIELD) and behavioral flexibility (STARR maze). B) for a 12-hole maze, showing maze rotation and shuttle replacement. 10-hole protocol is available in supplement. Turning of the maze prevents residual odors from providing predictive signals. The platform positions listed provide an example that does not reuse positions at the target during a specific training day. For targets, assure all four used targets are centered within a quadrant. Counterbalance across two cohorts to control for spatial bias.

The plastic platform was fixed firmly on top of a clean, white, plastic, 55-gallon container (33” tall, 26.5” wide (FG265500WHT, Grainger)). Although not required, a smoothly rotating maze with fixed locations will provide a much more reproducible maze set up.

At the minimum, locations should be taped/marked on the floor and the stand to allow reproducible turning of the maze (Fig. 1). In our setup, the 55-gallon container was secured to a RAM-PRO 12-Inch Heavy Duty Rotating Swivel Turntable/Lazy Susan (Amazon). The turntable was then fixed onto a 48” diameter wood tabletop (Menards). A sliding lock was bolted onto the white container such that it could be locked into any of the 10 (and later 12) positions. Care should be taken to align and fix (e.g., Loctite® Instant Mix Epoxy) the turntable, container, and maze-platform in a position such that the maze turns symmetrically and remains centered in the overhead video. Otherwise, software-based analysis will be more difficult, as the hole will shift slightly each time the maze is rotated. While some laboratories move external visual cues instead of rotating the maze, our preference is to allow multimodal spatial learning (sound, vibration, external odors, light gradients, etc.) of the external spatial environment for a more naturalistic task, quicker acquisition, and less exclusive reliance on vision. Whichever cues are used (all spatial modalities vs. only visual or only odor) should be deliberately chosen and reported, as they will likely impact outcomes.

### Room and platform setup

The testing area should be quiet and well lit. Our “room” was square, with walls ∼1 meter from the edge of the platform. The walls included 1) Windows with a blackout curtain (to prevent variability from sunlight) and visually distinct posters 2) a plain wall with a different poster, 3) a wall with shelves, and 4) a black room divider curtain (Proman Products Room Divider, Black [SKU: B07VJXTK87]). Smaller items in the room were marked to keep them in reproducible locations. A monitoring area near the testing area, but behind the curtain prevents interference by movements of the experimenter, with the experimenter always remaining in the same location behind the curtain. Flood lights (4 in total) were purchased from Amazon (Neewer Dimmable Bi-Color LED [Amazon]) and two each placed on opposing walls. Targets were always counterbalanced across groups to mitigate preferences for specific directions in the room. As some frame rates on some cameras match the flicker from flood lights, this should be checked carefully for compatibility.

Although the maze is rotated every trial, odor cues from prior trials can theoretically distract the mice, so we use 70% ethanol to clean the maze between trials. Shuttles and other items should be cleaned with an appropriate cleaning agent approved by local animal care guidelines. PLA plastic did not deform when cleaned with Alconox, 70% ethanol, or a peroxide--based cleaner (Rescue). We used 1% Alconox and water rinse, followed by air drying, to wash shuttles, moon caps, grates, and tunnels. An additional step of rescue cleaner was used between groups of mice in different cages, followed by 70% ethanol (maze) or 1% alconox (shuttles) for consistency.

The maze is set up by covering all decoy shuttles with grates, placing shuttles under every open hole, placing doors on the shuttles, sliding the shuttles underneath all active (target and decoy) holes, and covering the holes with moon caps blocking the view from the center of the maze (Fig 1). We recommend using an opaque/frosted release funnel (18cm diameter, 20cm height), with a plastic platform (can be 3D printed) to place the mouse atop the maze. The platform can be slid from under the mouse for a gentle release. Finally, a cart will be needed to sit the home cage on during shuttle docking, sized appropriately so the tunnel can reach and allow the mouse entrance into the cage. Covering the home cage with a drape (dark, covered space) after attaching the docking tunnel to the shuttle at the end of a trial facilitates mouse entry into their home-cage.

### Moon caps

Initially, we developed fully covered holes that required the mice to physically push off the cap to uncover the escape hole, which helped reduce serial checking. However, in early testing with a different 5xFAD cohort, we found that although all mice could learn to push a cap off the hole, some 5xFAD mice would “forget” how to push the cap by the following day, halting further trials (data not shown). This deficit prevented the further pursuit of fully covered holes; acquisition of “pushing caps” required two days of training, but if not retained, provided no advantage. This led to development of moon caps. All mice easily adapt to the optimized moon-shaped caps, which block the view of hole openings from the center of the maze but require less learning.

### Barries for the STARR maze protocol

For the STARR maze protocol (4^th^ day), the room and maze are identical to that described for the Barnes FIELD above except clear plastic barriers are added to create a clear radial-arm-style maze between each hole.

The barriers encourages the mouse to return to the center after each hole is checked. Barriers were clear, 0.250-inch-thick acrylic,-16“ X 3.5” thick (Professional Plastics, Sacr.250CCP), supported by 3D-printed barrier holders that slide onto a maze 0.635 cm thick.

### Mice

All experimental procedures were performed in accordance with relevant guidelines by the University of Iowa Institutional Animal Care and Use Committee (IACUC). Mice were housed on a 12:12 hour light:dark cycle and tested during the light phase (ZT+2 to ZT11). Experimenters tried to maintain consistent timing within groups (early or late in the light phase) but due to space constraints, studies often spanned across the day, which likely contributes to variability.

For experiments designed to test the shuttles, male and female C57BL/6J (#000664) were purchased from the Jackson Laboratory at 3 months of age and allowed to habituate to the animal colony room for at least 1 week following arrival.

### Procedure

#### The 4-day Barnes FIELD protocol

The protocol consists of four days of training and testing, as well as four separate probe trials to evaluate memory at different points in learning (early and late), as well as lengths of memory retention (short and long). A summary protocol for a 12-hole version is shown in Fig. 2, and a 10-hole version can be found in Supplemental Figure 6. A more detailed protocol is included in Supplemental Methods: “Barnes FIELD Procedure: Step-by-step”. Briefly, Day 1 starts with one habituation trial during which the bedding from the home-cage is placed in the escape shuttle to encourage the mouse to exit the maze without interference by the examiner. During the habituation trial, the shuttle should be placed in a location distinct from the training location on subsequent trials.

Other protocols commonly encourage pushing the mouse into the dark box if they require longer than 3 minutes to find the exit. By using this home cage bedding and connection to the home cage, our protocol generally eliminates the need for a negative reinforcer from the investigator, which may increase variability when used in only a subset of mice. The Day 1 habituation trial is followed by five consecutive learning trials with 3-minute inter-trial intervals. Day 2 starts with a probe trial (24h retention, early learning) followed by six more learning trials. Day 3 starts with another probe (24h retention, late learning), followed by six more learning trials and ending with one short-term memory probe trial (3-min retention, late learning).

A week later (“Day 4” of the protocol), starts with a single probe trial (1-week retention, late learning). We used day 11, so that the maze could be run Monday-Tuesday-Wednesday and the following Thursday each week (Fig. 2). This allows new cohorts of mice to be run every week without overlap. This one-week probe trial can be followed immediately by the STARR maze protocol detailed below to test behavioral flexibility. All trials have an inter-trial interval of three minutes.

We recommend that the time limit for learning trials be set high to improve analysis, as some outcome measures (distance, total visits, and unique visits) cannot be used if the mouse does not go to the target in a statistically valid manner. Some severely impaired mice require adaptation and there is the potential for confound regardless, but this can be mitigated with careful planning (see: Supplemental Methods: Troubleshooting, and Sup Fig. 3: Intervention protocol). Probe trials are conducted with all escape shuttles blocked by plastic grates and with duration limited to 90 seconds except as noted. A step-by-step protocol is included in Supplemental methods.

**Figure 3:**
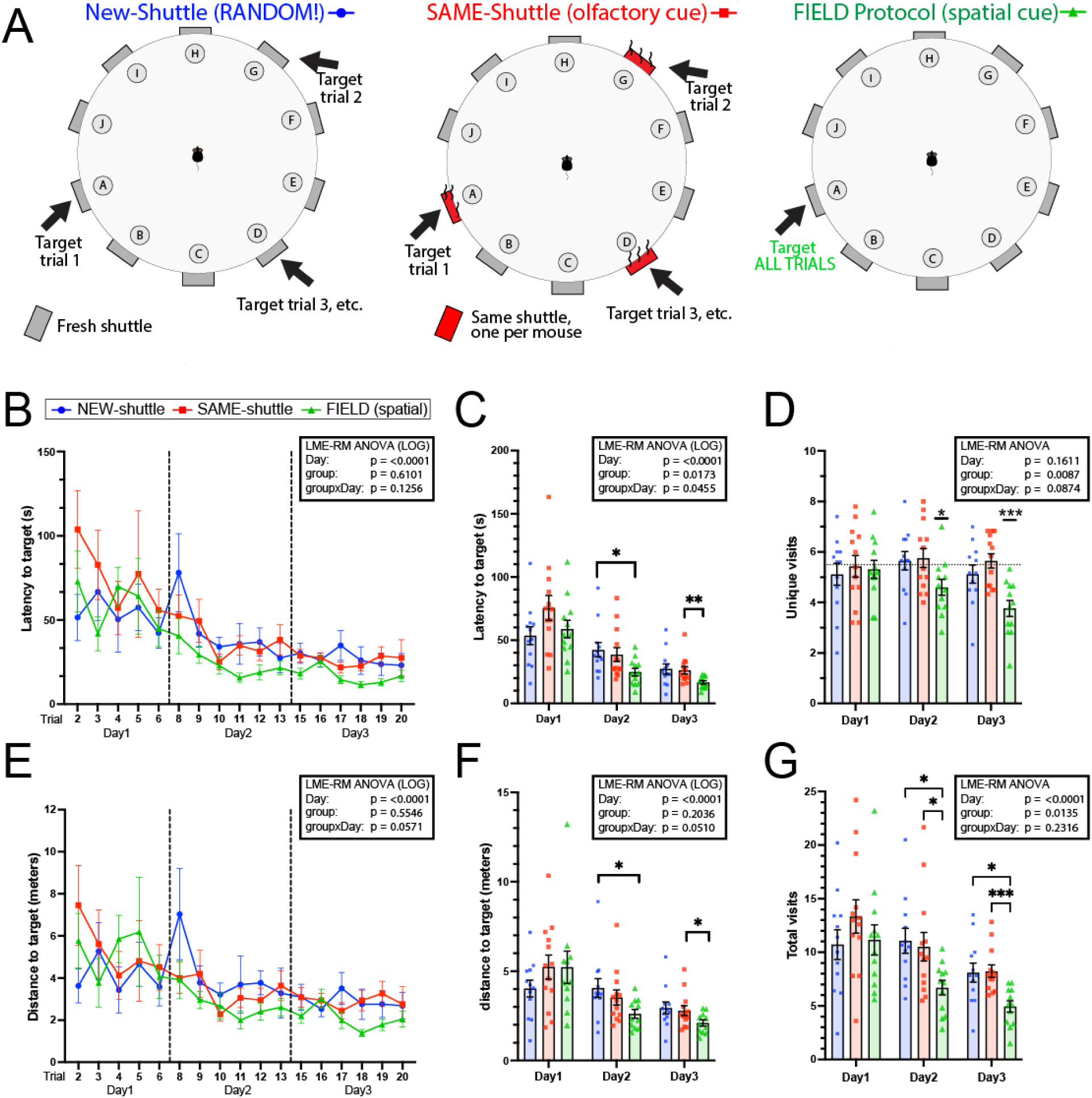
Total Learning vs. Spatial learning: Experimental design (A): Random (no-cue), SAME (uncleaned-shuttle as olfactory cue) and FIELD (normal spatial cue). Latency by trial (B) and day (C) reveal significant improvement across time in all three groups. Differences between groups are statistically evident when averaging by day, but the smaller magnitude of the difference demonstrates that <u>non-spatial learning</u> is a significant component of latency (B-C), distance (E-F), and total visits (G) outcomes. Unique visits (D) allow comparison against theoretical chance. Stars indicate days for which the group statistically outperformed chance (5.5 holes, one-sample Wilcoxon), only significant in the spatial group. The outcome “total visits” (G) includes repeat visits. N = 12 (NEW), 13 (SAME), 12 (FIELD). Linear Mixed-Effects Model Repeated Measure (LME-RM) using raw outcomes, or where indicated, natural log (LOG). Tukey correction for multiple comparisons. For clarity, only between-group comparisons shown. *p<0.05, **p<0.01 ***p<0.001

#### Spatial Training and Rapid Reversal (STARR) maze Protocol

Immediately following the standard one-week probe trial (Day 4, trial 1), barriers are added to the open platform to create a clear, radial-arm-style maze (Fig. 1, Fig. 8J), which is then used to test rapid and flexible learning of two new spatial target locations. A detailed protocol is included in Supplemental Methods: STARR maze Procedure: Step-by-step.

Briefly, the target for Trials 2-5 and Trials 6-9 are each in a different quadrant, distinct from the spatial location trained on Days 1-3. Eight learning trials are run with the barriers, with the same 3-min inter-trial interval used during the standard protocol. The last trial (Trial 10) is run as a probe trial, but with barriers. This final probe trial is evaluated in two ways. The first 90 seconds are analyzed to evaluate spatial preference, as with prior probes. However, the mouse is allowed to continue to explore until it has explored the rest of the maze (all but one hole). This task is sensitive to motivation, working memory and exploration strategy.

We do not recommend running the STARR maze protocol without first performing Days 1-3 of the standard Barnes FIELD protocol. Days 1-3 of the protocol provides information about spatial acquisition and long-term memory/retention, which may impact other tasks. Furthermore, since performance during early trials of any maze are highly variable, it is better to separate initial training from behavioral flexibility. Specifically, when we performed a single-day STARR maze protocol with no prior training, all mice completed the task, but more mice had difficulty acquiring the spatial location (Supplemental Fig. 4C + 4D). This is not surprising given most mice do not show robust evidence of spatial acquisition on the probe trials until completing two days (∼11 trials) of training. We also recommend against delays longer than a week between Days 1–3 and the STARR maze protocol because a cohort of mice tested after a month-long delay (Fig. 8B, 5xFAD littermate control mice), exhibited more difficulty than other cohorts. If the STARR maze protocol is used alone, delayed, or in mice with severe amnestic phenotype, we recommend at least one trial without the radial-arm barriers as habituation, prior to starting. Once acquired, the STARR maze can be repeated multiple consecutive days with new spatial locations. However, it should be noted that in control C57Bl6 mice, past 10 consecutive days, performance may decrease, likely as exploration increases (Data not shown).

#### Data Analysis

Data can be analyzed from videos manually or with computer software. Each outcome measure is discussed below. The investigator analyzing the data should be blind to treatment condition. In this study, we used AnyMaze (Ver. 6.36 and 7.48), EthoVision (Ver. 14 and 17), and cross-validated all studies with full or spot manual-scoring. A “visit” zone and “entry” zone are defined around each target to allow the software (or manual scorer) to know when the mouse has found the correct hole or exited the maze. For evaluating probe trials, we divide the maze into equal slices, such that one hole is within the center of each slice. For evaluating working memory during the STARR maze probe, an outer rim (∼56%) and center region (∼44%), are demarcated in the software where the barriers begin, such that entry to each arm can be recorded. In our maze, visits were counted if the mouse’s head approached the hole from the outside (open) edge, within 5 mm.

#### Outcome Measures: Considerations

Learning trials can be evaluated by different measures of efficiency (latency, distance, unique visits or total visits), while probe trials rely on measures of preference (percent of time near target). The primary outcome measure for each type of trial should be predetermined to allow accurate hypothesis testing.

Our general recommendation is to use <u>latency, averaged by day</u>, to evaluate general learning over Days 1-3 learning trials. Latency is often the most sensitive measure, encompassing multiple deficits. Unique visits can be used to check if findings specifically suggest deficits in the speed of <u>spatial acquisition</u>. Finally, we recommend percent time in <u>quadrants</u>, for probe trials, to test for spatial memory retention.

With new disease models, screening for a variety of deficits is sometimes desired, or the motivation behind a previously found deficit is not yet known. In this case, it may be warranted to compare additional outcomes. For example, a large difference in latency during learning trials with no corresponding difference in distance could suggest an arousal or anxiety deficit (periods of freezing or sleeping) or a motor deficit (decreased mean speed; Supplemental Fig. 4A and 4B). If multiple similar measures are evaluated, the cost of a type 1 vs. type 2 error should be considered.

**Figure 4:**
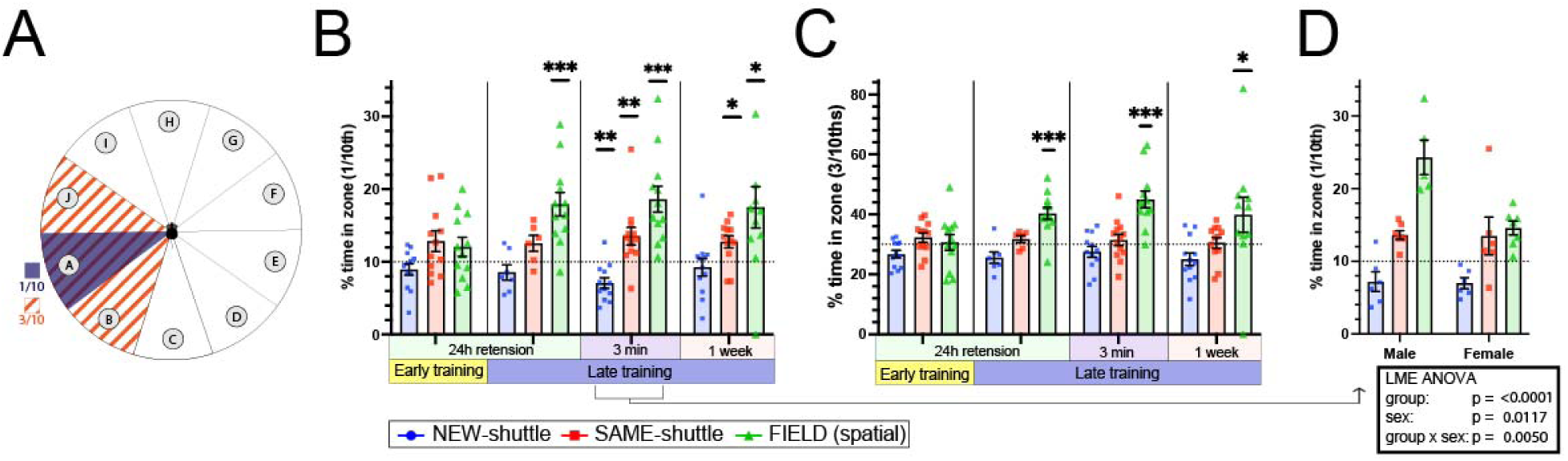
Intermittent probe trials for spatial acquisition and retention. (A) Schematic of the more precise 1/10^th^ versus traditional “quadrants” (here 3/10ths) of the maze to evaluate spatial memory. The odor cue will only be present in the 1/10^th^ sector (B) The percentage of time spent in 1/10^th^ of the maze following 24-hour retention at beginning of Day 2 (after 5 learning trials), and Day 3 (after 11 learning trials) and short-term memory at end of Day 3 (after 17 learning trials), and one-week retention. (C) The percentage of time spent in 3/10ths. (D) Comparing percent time (1/10^th^) between males and females on the 3-min retention trial. N = 12 (NEW), 12 (SAME), 12 (FIELD). One-sample Wilcoxon-signed rank test compared with theoretical mean of 10% (B) or 30% (C). (D) Linear Mixed-Effects Model Repeated Measure (LME-RM ANOVA), with p values corrected for multiple comparisons (Tukey) *p<0.05, **p<0.01, ***p<0.001, ****p<0.0001. 0.80 power, 0.05 alpha

**Figure 5:**
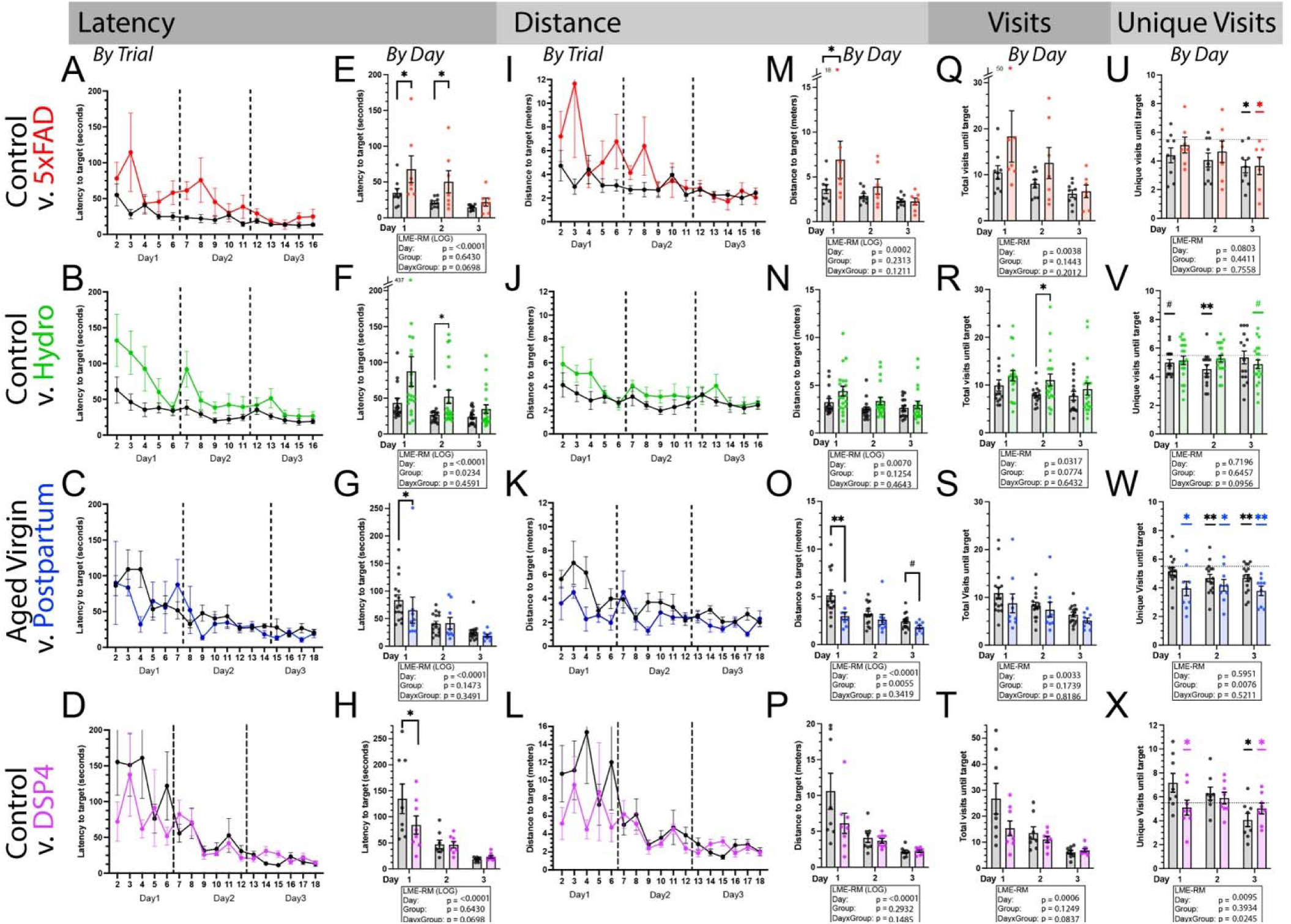
Outcome measures on learning trials across four models: 5xFAD (9-10 months-old, red), hydrocephalus (8-months-old, green), aged postpartum females (1 year, blue), and DSP4-injected (6-months-old, purple) compared with age-matched and/or control injection controls (black). **Latency** by trial (A-D) for visualization (no stats). Latency by day (E-H) shows significant main effect of day (learning) across conditions and experiments. All experiments also show differences in latency between groups on pre-planned comparisons, often on the first or second day. **Distance** by trial (A-D) for visualization (no stats); distance by day (M-P) shows a very similar learning curve as latency, but may reduce contribution of deficits in speed or immobility. **Total visits** by day (Q-T) is often similar to distance and can be tracked by hand if needed. **Unique visits** (U-X) allows comparison to chance. Mice with <u>no memory</u> of the cue on a 10-hole maze should <u>average 5.5 unique visits</u>, even with serial checking. A group average below chance for unique visits provides evidence for spatial acquisition (U-X). **Stats:** Linear Mixed-Effects Model Repeated Measure (LME-RM) ANOVA of raw or natural logs (LOG). For unique visits, each probe was separately compared versus chance (10 holes: 5.5 (U,V,W), 12 holes: 6.5 (X)) using one-sample Wilcoxon sign-ranked. p*<0.05 **p<0.01. Non-significant trends (#p<0.1) are shown for illustrative purposes.

### Learning Stage (Learning Trials)

#### LATENCY

This is the time (seconds) it takes the mouse to find the target hole. Latency is sensitive to confusion, time spent orienting, planning, freezing, grooming, sleeping, search strategy, motor deficits, overall speed, and motivation.

#### DISTANCE

This is the distance (meters) it takes the mouse to find the target hole. It is less impacted by motor deficits and penalizes mice that follow the edge (thigmotaxis), even if they remember the spatial location. It is insensitive to freezing, orienting, sleeping, or planning (i.e., if the mouse isn’t moving). Distance may be an appropriate first choice measure if the model has known motor deficits, but the hypothesis targets cognitive changes.

#### UNIQUE VISITS

The number of holes checked once to find the target hole. The range is 1-10 for a 10-hole maze. Statistical chance is 5.5 holes visited for a 10-hole maze, or 6.5 for a 12-hole maze (expected = (N+1)/2). When averaged across trials (one animal) or across animals (one time-point), this outcome measure allows us to test whether a mouse or group outperformed chance. It is the only learning trial outcome measure that can provide direct evidence that a specific association (e.g. spatial location, odor cue) has been acquired.

#### TOTAL VISITS

The total number of holes checked, including repeats, (range 1–unlimited) to find the target hole. Statistical chance (assuming a mouse had no working memory and no ability to perform serial checking) would average 10 total visits for a 10-hole maze. While finding a real model that has “zero working memory” is unlikely, it can help contextualize the findings. For example, a mouse that consistently returned >10 times to the same arm would be demonstrating abnormal ***perseveration***, not impaired memory, as this many incorrect visits would be inconsistent with chance alone.

#### DAY-BY-DAY outcomes

Latency, distance, unique visits, and total visits can be averaged across trials within the same day. We recommend this as the primary outcome measure due to its straightforward statistical analysis and reduced noise. Visually examining the trial-by-trial data may still be of interest as the highest rate of learning is usually within the first several trials and differences between groups may be obscured by averaging if learning rate is the main deficit.

#### TRIAL-BY-TRIAL outcomes

All outcomes can be examined by trial, but are highly variable due to the chance component inherent in the Barnes maze; even the worst-performing mouse has a 1/10^th^ chance of hitting jackpot on each trial. Furthermore, statistical tests must be chosen carefully when analyzing across many repeated trials (see statistics). If trial-by-trial differences are planned as the main outcome because changes are expected only in early trials, a larger sample size will be beneficial to mitigate the effect of chance. This is expected variability when evaluating early learning rather than an over-trained outcome.

To reduce random noise for illustrative purposes, we show group averages for trial-by-trial graphs and individual mouse data points for day-by-day graphs (Fig. 3).

#### Probe trial outcomes: Memory and Motivation

Our Barnes FIELD protocol includes four separate memory probes to test for evidence of **spatial learning and retention,** independent of other **general learning mechanisms** (e.g. meta-learning, motivational changes, procedural learning and habituation) that dominate gains in learning trials. Three outcome measures on probe trials were evaluated:

#### Percent Time in Quadrant

Quadrants (or equivalent) are the most common outcome used for both the Barnes maze and the Morris Water Maze. For a 12-hole maze, a quadrant is the target hole and two adjacent holes (3 of 12), representing 25%, of the maze (Fig 6A & 6B). For a 10-hole maze, 30% of the maze, consisting of the target hole and two adjacent (3 of 10), is the best approximation, as mice tend to examine the target and nearby holes. Time spent is compared to the actual percent of the maze expected by chance.

**Figure 6:**
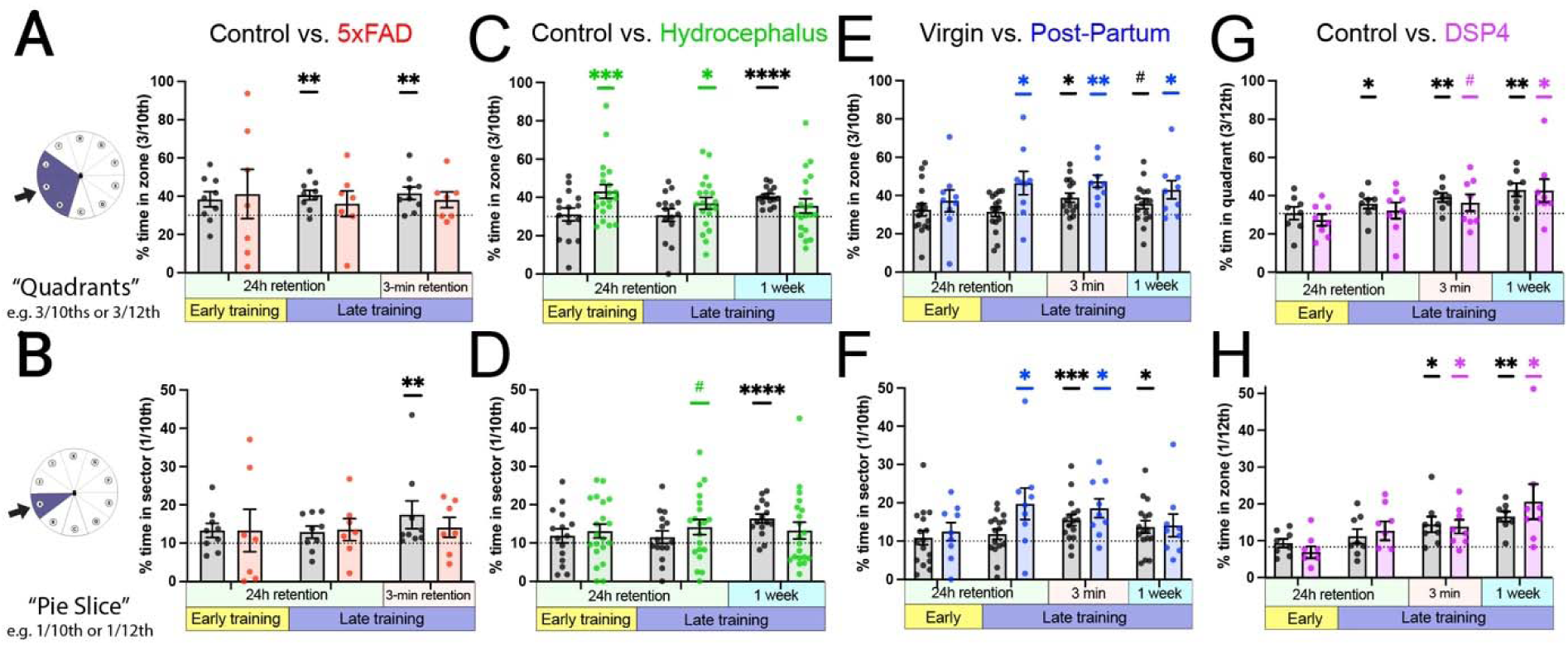
**Probe trials provide statistical evidence of spatial acquisition**. (A) 5xFAD mice as a group fail to show evidence of spatial learning compared to chance, while age-matched controls show evidence in late training. (B) Controls also show evidence when a more precise sector is analyzed at the 3-min retention probe. (Control: n = 9, 5xFAD: n = 7, no one-week probe). (C) Controls in the hydrocephalus experiment only show evidence of spatial acquisition at one-week (no 3-min probe performed), whereas the hydrocephalus manipulation shows evidence on Day 1.(D) (Control: n = 15, Hydrocephalus: n = 20). (E) Postpartum mice show evidence of spatial acquisition in all probes following the first day, whereas aged virgin mice require longer training. (F) Results are relatively similar for the more precise sector. (Virgin n = 16, Postpartum: n = 9). (G) Control mice show evidence of spatial acquisition in all probes following the first day, whereas DSP4 require more trials to outperform chance. (H) Results are relatively similar for the more precise sector, but might alter interpretation (Control n = 8, DSP4 n = 8). STATS: One-sample Wilcoxon signed-rank against theoretical mean of 30% (A, C, E) and 10% (B, D, F) for 10-hole and 25% (G) and 8.3% (H) for 12-hole maze. p*<0.05. **p<0.01, ***p<0.001, ****p<0.0001. Non-significant trends (#p<0.1) are shown for illustrative purposes.

**Figure 7:**
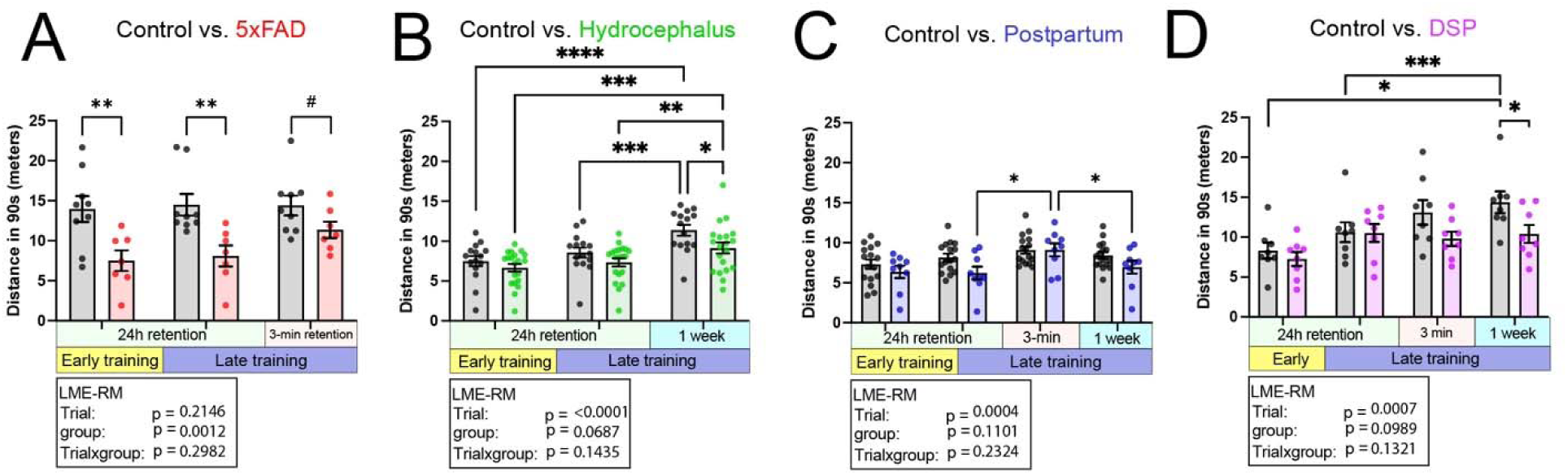
Total distance travelled across 90-second probe trials is highly sensitive to motivation and behavior. (A) 5xFAD mice show significantly reduced distance travelled across probe trials (main effect genotype) (n = 9,7). (B) Hydrocephalus model shows increased distance traveled from Day 1 to Day 4 in both groups of mice, but the control mice travel significantly more on the one-week probe trial than the Kaolin-injected mice (main effect of trial and group) (n=15, 20). (C) For the postpartum experiment, there is a main effect of trial, but no interaction (n=16, 9). (D) The DSP4-injected mice do not increase distance traveled at 1 week, whereas the control mice show increased distance from Day 1 to Day 4, and travelled more on Day 4 than the DSP4-injected mice (n=8,8). Linear Mixed-Effects Model Repeated Measure (LME-RM) ANOVA, with Tukey for multiple comparisons. p*<0.05. **p<0.01, ***p<0.001, ****p<0.0001, Non-significant trends (#p<0.1) are shown for illustrative purposes.

#### Percent time at Sector

The percentage of time investigating the more precise pie slice (sector) containing only the target (e.g. 1/10^th^ or 1/12^th^). Using a precise sector is needed when purposely training with non-spatial cues (e.g., odor; Fig. 4). Our recommendation is to evaluate “quadrants” unless testing a non-spatial cue or if spatial precision itself, rather than spatial memory retention, might be impacted by the manipulation. In either case, time spent above chance suggests evidence of memory, while excessive time at that location may suggest perseveration.

#### Total distance travelled

Total distance during the 90-second probe trial encapsulates the speed and motivation of the mouse, including hyperactivity, speed, and perseverance. An ideal strategy is to check the expected hole, and once the expected hole is found to be blocked, explore the rest of the maze for alternatives. A perseverative mouse might stay at the expected hole and not move on, whereas a non-motivated or fearful mouse might fail to explore at all. Hyperactivity relative to control animals would include travelling a greater distance, with or without efficient exploration of all sectors of the maze.

### STARR maze OUTCOMES

The STARR maze is identical to the Barnes FIELD protocol except for the addition of radial-arm barriers used to reduce serial checking. Additionally, the mouse is expected to learn two new locations rapidly to test flexible acquisition. The rapid reversal can also be trained without barriers, and is comparatively easier. We have published studies with the non-barrier rapid reversal protocol previously [30].

### Rapid Reversal Learning Trials

While all of the same outcome measures could be used (latency, distance, unique visits, total visits), based on our findings in the Barnes day 1-3, we chose latency to evaluate all learning, and “unique” to test for evidence of spatial acquisition. Some experimental groups show a large increase in latency when the spatial target is switched, providing evidence they recognize the location has moved. By contrast, the unique visits outcome provides statistical evidence that the new location is being acquired trial-by-trial.

### Final Probe trial

#### Percent time in each sector (90-second probe)

Relative balance of time spent at most recent (reverse target), relatively recent (new target), and originally-learned location (Barnes target Day 1-3) provides information about recent and remote memory preference, but should be interpreted cautiously and in the context of prior performance.

#### Repeat Entries (extended probe)

The cumulative number of “errors” (repeat entries) made while exploring a set percentage of the maze (N-1 holes). The probe is extended in time until all but one hole has been visited. (If mice do not visit all holes promptly, see Supplemental Fig. 3). Repeat entries on a radial arm maze or during a spontaneous alternating task are often used as a measure of working memory errors [31]. However, repeat entries should be interpreted carefully, as increased repeat entries may also be evidence of a win-stay strategy (preference and memory for prior location), and decreased repeat entries may be evidence of serial checking or tendency towards win-shift or exploration strategy.

#### Time for complete exploration (extended probe)

The time needed for the mouse to explore the majority (N-1 holes) of the maze is sensitive to motivation, strategy and motor deficits).

#### Quantitative analysis with automated software

Latency, distance, and total visits are all available via AnyMaze. Unique visits, time in quadrant, and time in sector can be derived from these using Excel formulas or via custom-MATLAB scripts. Total visits extracted directly from AnyMaze encompasses any approach to the target (including “double takes”). For repeat visit calculations, we only count repeat visits that are separated by a check to another arm. For this, we created a separate MATLAB script, which only counts sequences that include the center and another arm.

### Statistics

#### Data normalization

Evaluation of distance and latency suggested these outcomes were non-normal, particularly in more impaired mice. As these outliers likely represent true deficits, rather than equipment error, we normalized our data. Following analysis of experiment 1 (olfactory experiment), we determined that taking the natural log of these right-skewed outcome measures resulted in more normalized data, allowing a standardized approach across experiments without need to exclude outliers. Natural log data was used for all statistical comparisons for latency and distance, but raw data was graphed to allow visualization of expected outcomes in real terms. Unique and total visits did not require transformation.

### Hypothesis testing and choice of outcome measures

#### Daily Averages

We used a repeated measures mixed effects model (REML), with ANOVA output table in Prism (GraphPad, ver10) for all daily averages, including distance (natural log), latency (natural log), total visits, and unique visits for all experiments. In most cases, a standard repeated measures ANOVA is equivalent. When the number of repeated measures is low (e.g. less than 5), a general linear mixed-effects model from any statistical package should also be appropriate and generally equivalent. For Fig. 3, we also compared prism output to R and MATLAB and found results to be equivalent. Sphericity should not be assumed, as early trials and days are usually more variable.

#### Trial-by-trial

We do not recommend evaluation by trial, except in specific cases, because of the high number of repeated time points. Using standard repeated measures ANOVA with > 6 time points, especially with small sample sizes, may increase the risk of violating sphericity and can inflate type 1 error [32]. If it is essential to evaluate trial by trial changes, evaluation separately within each day is one possible method. Alternatively, modeling with a more robust mixed-effects model may be utilized. Statistical consultation is recommended if evaluation across all trials together is desired.

In our study comparing these outcome measure, trial-by-trial data for Says 1-3 (Fig. 3) was analyzed using a linear mixed-effect model in MATLAB:

lme_all = fitlme(learntbl,’LogDistance ∼ group*ConsTrial + (1|Animal)’,’DummyVarCoding’,’effects’,’FitMethod’,’REML’);

The listed p-values are derived with an ANOVA-style table for main effects and interactions. The results were checked separately by our statisticians with R, using an equivalent model. Unlike data averaged by day or examined within day, statistical p-values differed more when analyzed across all trials using different statistical packages, leading to the cautions listed above. However, conclusions were largely equivalent.

#### Planned Comparisons

Comparisons were made between groups at each averaged day for latency, distance, and total visits, and corrected using Tukey in Prism when indicated. For clarity, paired comparisons across time were often omitted from graphs, even when significant.

#### Comparison to chance

For average unique visits (during learning) and percent time spent (during probe trials), each group’s performance was tested against statistical chance using a one-sample Wilcoxon sign-rank test. A one-sample t-test would also be appropriate in most cases, but we chose a non-parametric test for consistency across experiments. We do not suggest correction for these pre-planned tests, as this would inflate type 2 error and may erroneously miss when groups begin to show evidence of spatial retention. The risk of type 1 error should be mitigated by interpreting within the context of collaborating evidence.

#### Power and sample size

Our primary experiment compared spatial cues (normal Barnes) vs. random chance (Fig. 3), thereby allowing us to estimate a minimum hypothetical sample size necessary to detect differences attributable to <u>spatial learning</u> for each possible outcome measure (Table 1). Sample size was calculated using MATLAB using a power of 80%, and alpha of 5%, with the following:

**Table 1:** Sample size based on calculations from mice with spatial cue compared with no cue (Fig. 3+4): To detect differences on learning trials due to deficits specific to spatial learning, we recommend a minimum sample size of **15 per group** (average sample sizes across outcome measures day 2 + 3) for latency. For detection of evidence of learning from probe trials, as few as **5 subjects** may be sufficient for a 3-min delay, but would require complete loss of spatial acquisition in one group.

| **Learning Trials** | **Day 1** | **Day 2** | **Day 3** |  |  |
| --- | --- | --- | --- | --- | --- |
| Latency | 211 | 12 | 19 |  |  |
| Distance | 87 | 16 | 23 |  |  |
| Unique | 737 | 21 | 13 |  |  |
| Total Visits | 1952 | 10 | 11 |  |  |
| **Probe Trials** |  | **Day 2-**<br>**24h** | **Day 3-**<br>**24h** | **Day 3-**<br>**3min** | **Day4- 1**<br>**week** |
| Probe (spatial vs.<br>Rand) |  | 49 | 4 | 5 | 14 |
| Probe (spatial vs.<br>chance) |  | 1739 | 4 | 4 | 32 |

n = sampsizepwr(“t2”, [m1, pooledStd], m2, 0.9, []).

Sample size needed to detect differences due to non-spatial learning or other group differences will vary depending on the effect size.

#### Mouse models

We sought to test our protocol as a screening tool for spatial and non-spatial learning, spatial memory, motivation and behavioral flexibility across several mouse models of cognitive change. Studies were performed by laboratories at the University of Iowa who requested assistance with cognitive evaluation of mice between 2020-2025. Each laboratory was taught the current protocol and used the same setup, making minor changes if needed for their experimental constraints. Importantly, teaching the task required only 1-2 hours and the investigating laboratory performed it independently (in most cases). Data presented here were reanalyzed with permission and include data from mice used in previously published work.)

#### Aged Postpartum

∼12 month-old C57BL/6J females (n=25) with and without a history of pregnancy were administered sertraline (167mg/L oral suspension, *ad libitum* via in-cage water bottles for ∼8 weeks) or water [27]. For our analysis, groups were collapsed across treatment to focus on comparing outcome measures, using only the postpartum vs. virgin condition as the experimental manipulation.

## 5xFAD

8- to 9-month-old male (n=7) and female (n=9) heterozygous 5xFAD (n=7) vs. littermate controls (n=9, Tg(APPSwFlLon,PSEN1 *M146L*L286V)6799Vas, Jackson Laboratories). Only non-drug-treated mice were analyzed for this methods paper [25].

### Kaolin injection as a model of Normal Pressure Hydrocephalus (NPH)

8-month-old C57BL/6J mice (N=35, 19 female, 16 male) received a 5 µL injection of either sterile saline (n=15) or a 15% suspension of kaolin to induce hydrocephalus as a model of normal-pressure hydrocephalus (NPH; n=20) [26].

### DSP4 toxin

DSP-4 (N-(2-chloroethyl)-N-ethyl-2-bromobenzylamine hydrochloride) is a neurotoxin thought to impact nerve terminals of noradrenaline neurons in the locus coeruleus and brain regions innervated by the locus coeruleus [33–36].

The locus coeruleus shows pathology early in Parkinson’s disease and Lewy Body Dementia [37–39], diseases characterized by non-amnestic cognitive impairments. 3-4 month-old C57BL/6J mice (*N* = 16, 11 females, 5 males) received intraperitoneal injection of either DSP-4 (50 mg/kg, Sigma-Aldrich) dissolved in saline (*n* = 8) or saline alone (*n* = 8). Mice were injected twice, 7 days apart [40–42]. The Barnes FIELD and STARR maze protocol began 4-7 days after the second injection.

### Olfactory Shuttle Experiment

3-month-old male and female wildtype (WT) C57B6J mice (n= 37) were randomized and interleaved into three separate conditions (Fig. 3A). In the first group, “NEW-shuttle,” (N=12, 6 females, 6 males) used shuttles were exchanged for clean shuttles every trial, and the spatial target for each trial was pseudo-randomly changed. By design, any improvements across trials should reflect non-spatial learning (e.g. meta-learning, search efficiency, velocity, motivation, etc.) as the random location should theoretically make all potential cues non-predictive. In the second protocol, “SAME-shuttle,” (N=13, 6 females, 7 males) the same target shuttle was used for all trials across all four days.

The shuttle was not cleaned for olfactory cues between trials, but each mouse had its own personal shuttle copy. As with group 1, the target spatial location was changed randomly for each trial. The third group ran our proposed Barnes protocol with standard spatial cues, the “Barnes FIELD” protocol (7 females, 5 males, *n* = 12), an optimized spatial training protocol including printable individual shuttles. In this protocol, escape shuttles that are entered are never used more than once per >10 trials (and always washed) and moon caps hide the entrance so that only external spatial cues are available until approaching the targets.

## RESULTS

Two obstacles limit estimating learning rate during task acquisition in land-based mazes: first, mice might use olfactory or intramaze visual cues to locate the target escape box during learning trials. Second, the literature is mixed as to which outcome measures (e.g., latency, distance, etc.) are most sensitive and specific to deficits in learning rate.

To mitigate these obstacles, we designed new escape boxes that could be printed in replicate. To determine whether use of the shuttles provides superior masking of potential cues compared with standard protocols, we tested whether mice with a reused escape shuttle (SAME-shuttle group) could utilize subtle olfactory cues in the absence of spatial information. We compared this group to our recommended protocol (Barnes FIELD) and a random design with no cues (NEW-shuttle). The experimental design of the olfactory shuttle experiment further allowed us to compare the sensitivity of several outcome measures in their ability to detect spatial compared with non-spatial learning.

### I: Distance and latency gains largely represent non-spatial learning, though the benefit of spatial cues is statistically detectable

#### Latency

The time, in seconds, to find the target (latency) is the most commonly reported outcome measure for learning trials [43]. All three groups showed significant changes in latency to find the target across individual trials (Fig. 3B, main effect: trial) and when compared across days (Fig. 3C, main effect: day). When comparing latency across all three groups trial-by-trial, the main effect of group was not significant (p=0.6101). However, the small benefit of spatial learning on latency was detectable in the day-averaged data (group x day interaction, p=0.0455). Pre-planned comparisons demonstrated a benefit for the Barnes FIELD protocol compared with NEW on Day 2 and OLD on Day 3 (Fig. 3C). There was no difference between the OLD and NEW shuttle group, suggesting that if mice use the olfactory cue to guide their search, it does not improve latency.

#### Distance

Distance to target removes influences attributable to motor speed, a potential factor for neurodegenerative models with motor features. However, examining only distance may reduce sensitivity to time spent orienting, freezing, planning, grooming or napping. In this experiment, distance to target across all groups also showed a significant main effect by trial (Fig. 3E, p=<0.0001) and by day (Fig. F, p=<0.0001), providing evidence for learning across time. Differences between the groups visually mirrored latency, suggesting using one measure over the other does not increase the sensitivity for spatial learning rate compared with non-spatial learning rate. However, latency reached significance for main effect by group, while distance was borderline (p=.05) for a group by day interaction.

### II: Unique visits provide statistical evidence for spatial learning

#### Unique Visits

While distance and latency are most classically used to evaluate learning trials, evaluating unique visits provides the opportunity to test whether the group is performing better than statistical chance during learning, rather than relying on preference during the probe trial. Statistically, a mouse with no available cues should, *when averaged over enough random trials*, visit 5.5 holes to find a target hidden amongst 10 identical holes. For each group, we tested individually whether the group “outperformed chance” on each day.

As hypothesized, the NEW-shuttle group did not differ from the theoretical mean of 5.5 on any of the three days (one-sample Wilcoxon rank sign test against hypothetical mean of 5.5), providing evidence that our 3D printed shuttles and moon caps successfully hid extraneous cues (odor or visual) during learning trials. Interestingly, the SAME-shuttle protocol also did not differ from chance across days, suggesting mice did not generally use the odor cue to gain advantage in reaching the target. By contrast, by Day 2 and 3, the FIELD-protocol group differ significantly from chance (Fig. 3D: Day 2: p=0.0166, Day 3: p=0.0005). This provides statistical evidence that by the second day of training, mice on a traditional protocol are using spatial cues to help navigate.

Furthermore, comparisons between groups using ANOVA show a significant main effect of group. Thus, spatial learning may be more easily detectable using unique visits as an outcome measure, while latency and distance provide estimates that include both non-spatial and spatial (or cue-based) learning.

#### Total visits

Finally, we examined the number of total visits, which includes both unique visits (as above), as well as repeat visits, which could be indicative of perseveration or working memory errors (Fig. 3G). Here, we found, much like distance, mice in all groups tend to complete the task with fewer visits to incorrect holes with each consecutive day (main effect day p<0.0001). There was also a significant main effect of group (p=0.0135), with pre-planned comparisons indicating larger improvements in the FIELD-protocol group, indicating benefit from use of the spatial cues on Days 2 and 3 compared with the NEW-shuttle and SAME-shuttle groups.

#### Power and sample size

We calculated the sample size needed to detect differences between the spatial (FIELD-protocol) and random (NEW-protocol) on each day, as this is useful as a starting point for experimental design when spatial learning is the outcome of interest (Table 1). Based on this experiment, around **15 mice per group** are needed to detect difference in spatial learning during the acquisition phase, as most gains are related to other forms of learning. The probe trials, by contrast, require as few as **5 subjects** to provide evidence for spatial acquisition.

### III: 3D printed shuttles mitigate the risk of olfactory cues in traditional protocols

Although the olfactory cue did not improve any outcome on the learning trials, we next sought to use the probe trial to determine if mice were still acquiring the olfactory cue from a reused shuttle. In probe trials, entry was blocked by plastic grates on each shuttle to prevent entry. Within SAME-shuttle group, the “reused” shuttle was placed in a new pseudo-random spatial location for the probe trial. We excluded mice whose shuttle box was randomly assigned to the location of the prior trial during the probe, to avoid spatial bias in the results. However, inclusion of these mice did not alter conclusions.

Intriguingly, we found that mice can identify the olfactory cue from a randomly placed reused escape shuttle. They learn and retain this cue over a similar timeframe as groups using spatial cues. At the short-term memory probe and the 1-week long-term probe, mice in the SAME-shuttle group outperformed chance, spending more than 10% of their time exploring the 1/10^th^ sector containing the target shuttle (Fig. 4B). By contrast, mice in the NEW-shuttle group did not spend a greater amount of time in a designated “target” sector.

Evidence for memory of the olfactory cue was only detected on probe trials when evaluating 1/10^th^ of the maze (sector with one hole). The mice did not beat statistical chance when evaluating the “quadrant” (3/10^th^s of the maze in this experiment). This may be due to the nature of the odor cue, which would only exist at one target hole. By contrast, with training using a spatial cue in the FIELD-protocol, there was evidence of increased time spent regardless of whether a more precise sector (1/10^th^) or broad “quadrant” (3/10^th^) was used (Fig. 4B and 4C). Use of a narrow vs. more spatially generalized portion of the maze for the probe trial may be sensitive to different aspects of cognition and was explored further in the mouse model case studies.

### IV: Outcome measures generally agree, but provide context

Our prior experience demonstrated that outcome measures do not always match; for example, latency to target can be increased while distance between groups is unchanged for a manipulation that impacts arousal (Sup Fig. 4A and 4B). For this reason, we decided to compare outcome measures across multiple cognitive models to help plan future studies and interpret existing data.

We evaluated latency (by trial and by day, Fig. 5A-H), distance (by trial and by day, Fig. 5I-P), and total visits by day (Fig. 5Q-T) in models performed by four separate laboratories for evidence of learning rate and differences between groups. Across most experiments and outcome measures, there was a significant main effect of day, demonstrating learning (spatial and total learning) across most models and controls.

Importantly, differences between groups were often evident only early in training. For example, 5xFAD mice had more difficulty finding the target, evident statistically on Day 1 for distance and latency, and Day 2 for latency (Fig. 5E). By Day 3, there are no differences between 5xFAD and littermate controls. This is confirmed by unique visits, which suggest both groups outperformed chance by Day 3, and thus had acquired the spatial location (Fig. 5U).

Within the NPH mouse model, latency (Fig. 5F) and total visits (Fig. 5R) showed differences at Day 2 between the groups, and all other outcomes showed the same trend. Unique visits (Fig. 5V) showed control mice were performing better than chance by the second day, while NPH mice did not. However, neither group outperformed chance on the third day. In all other groups, mice outperformed chance for unique visits during learning trials by the third day or sooner (Fig. 5U-X). The poor performance on Day 3 within the NPH experiment may represent realistic behavioral variability or a potential hidden factor.

Interestingly, aged postpartum mice learned very quickly within the first few trials of Day 1 compared with aged virgin female mice (Fig. 5M), outperforming chance as a group even on Day 1 (Fig. 5W). This finding is convergent with the trends on early probe trials suggesting early acquisition of the target by many of these mice (Fig. 6E).

In DSP4 injected mice, we hypothesized we would see no differences in learning rate on Barnes Day 1-3, but deficits in the flexible sessions in later stages of the STARR maze protocol. In agreement with this, learning rates and retention were relatively similar between the groups. Interestingly, as with the postpartum mice, some DSP4 mice showed an early advantage on Day 1. However, in this case the probe trial (Fig. 6G) did not support evidence that they had early spatial retention. Thus, these differences could have been driven by non-spatial learning, or since DSP4 mice also outperformed chance on unique trials, the information may have been acquired but not yet retained during the 24h delay.

Across experiments, evaluation of unique visits acts as a complement to latency/distance (discussed above) and probe trials (discussed below). Notably, the unique visits outcome agrees relatively reliably with the probe trials (Fig. 6), even though it is derived from <u>independent</u> trials. Probe trials are not included as part of the learning rate to avoid double representation of these trials in analysis.

### V: Probe trials evaluate spatial acquisition and retention across learning

Probe trials are used to test for spatial acquisition and retention. Probe trials include two 24-hour retention probes at the beginning of Day 2 (after 5 learning trials on Day 1) and Day 3 (after 11 cumulative learning trials). The 3-min retention and 1 week retention are both performed after the final learning trial (17 learning trials total). Not all of our model experiments included all possible probe trials (Fig. 6). We evaluated all groups with a more precise (1/10^th^) vs. broad (3/10^th^) spatial location to compare how this variable might impact results. In general, more groups across experiments showed the ability to outperform chance when using the broader “quadrant”, suggesting “quadrant” may be more sensitive to detecting evidence of spatial learning. A more precise sector, such as 1/10^th,^ may be more appropriate in certain situations (e.g. odor cue in Fig. 4B).

Importantly, the general trends remained similar across experiments regardless of whether a broad or narrow spatial location was used as the outcome.

### VI: Interleaving probe trials evaluates early and late acquisition, long-term (24h and 1 week) memory and short-term (3-min) memory

In a probe trial, all exits are closed. Time spent at the spatial target that is greater than would be expected by chance provides strong evidence of acquisition and retention.

Interleaving probe trials with learning trials allow testing acquisition early and late in training, and retention over short and long intervals. Analysis during acquisition is important as many animal models learn and retain the task, but at a slower rate.

Protocols sometimes omit probe trials over concern for extinction of learned spatial cues. Although we did not directly test learning with and without the probe trials head-to-head, we were usually able to detect evidence of spatial memory in control groups across probe trials (Fig. 4B-C, Fig. 6) despite including up to four separate probe trials. In our shuttle odor experiment, the FIELD group showed statistical evidence of memory retention at all probe trials except for the first day (Fig 4), demonstrating a typical time course of acquisition.

We did find that control mice in each study learned at different rates, likely due to age, background strain and other behavioral tasks being used by different laboratories. In the 5xFAD study, controls showed evidence of memory retention at the beginning and end of Day 3 (Fig. 6A). By contrast, the control mice in the NPH experiment only outperformed chance at the one-week probe (Fig. 6C).

### VII: Probe trials are subject to multiple motivational factors

Although the main outcome of the probe trials (percent time in quadrant or sector) is theoretically less influenced by motor speed compared with latency on learning trials, it is not immune to motivation and motor speed differences. A mouse that moves more quickly or is more efficient in exploration will take less time to explore a sector. This efficient mouse might spend relatively less time in the target sector over 90 seconds, despite adequate spatial acquisition. Evaluating other behavioral outcomes can help detect possible motivational or motor differences between groups. For example, we evaluated total distance travelled during the probe trial and found it was highly sensitive to differences between groups.

5xFAD mice showed significantly reduced total distance in 90 seconds across probe trials (main effect group, Fig. 7A). However, this effect was less notable in the 3-min probe, suggesting a potential interaction with motivation or memory, rather than a motor deficit.

**Figure 8:**
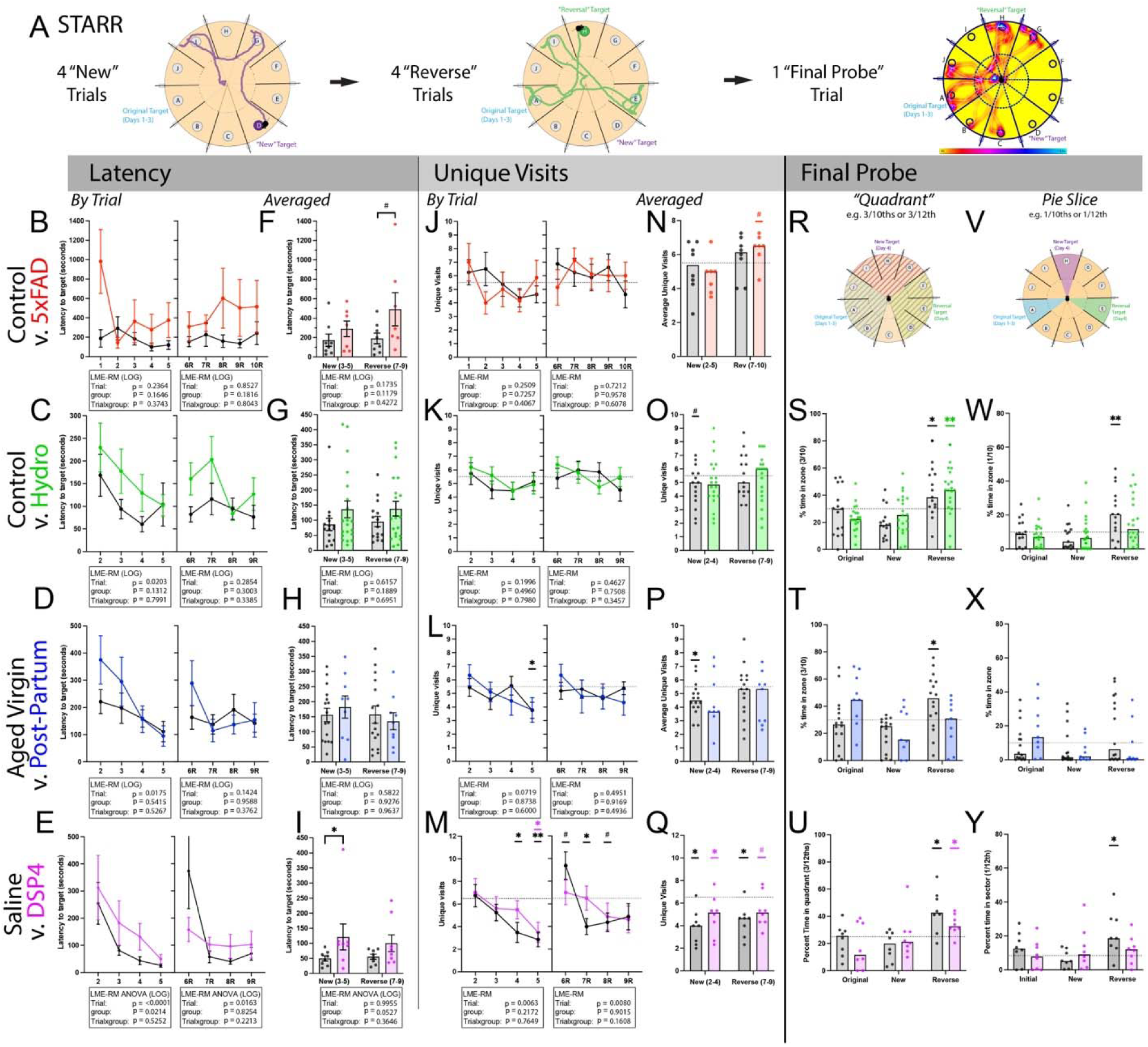
Spatial Training and Rapid Reversal (STARR maze): (A) Following 3 days learning one target during the Barnes FIELD, mice learn two new targets, with 4 trials for each. **Latency** (B-E) to find a new target shows improvement in most groups across new trials (2-5), but more difficulty with the final target (6R-9R). **Averaged latency** (F-I), excluding trials when targets change, allows evaluation of performance of individual mice. **Unique visits** allow comparison to chance (5.5 for J-L; 6.5 for M) at each trial or averaged by stage (N-Q). **PROBE trials** demonstrate preference/memory, with typical mice favoring the most recent (“reverse”) quadrant (R) and sector (V), despite difficulty “finding” this target. Statistical evaluation against chance was only performed for the reverse quadrant, but other regions are shown. Both groups in the hydrocephalus experiment show evidence of memory for the most recent target (S), though only the controls outperform chance at the precise zone (W). (T, X): Intriguingly, only non-postpartum mice favored the most recent target. (U,Y) Both groups in the DSP4 experiment showed evidence of memory for the most recent target (U), though again only the controls outperform chance at the precise zone (Y). p*<0.05. **p<0.01. All data are graphed as raw for illustration, stats use natural log (LOG) where indicated. Non-significant trends (#p<0.1) are shown for illustrative purposes.

Distance travelled in 90 seconds differed between probe trials during different stages of the experiment for both saline and kaolin injected mice (Fig. 7B) as well as postpartum (Fig. 7C) and control (non-DSP4) (Fig. 7D) groups. Since motor function should not differ between these days, these differences across time likely represent motivational changes, possibly with more exploration in later probe trials when the mouse is more confident in the location.

Finally, by inspection, we noted substantial differences between control groups across experiments. These differences may relate to genotype, strain, age, experimenter or any additional behavioral tests completed by each group. 5xFAD mice, with the highest distance travelled (Fig. 7A) are on a mixed background strain compared with the other experiments using C57BL6 (Fig. 7B-D). 5xFAD have prior reports of hyperactivity [44].

### VIII: The Spatial Training and Rapid Reversal (STARR) maze challenges behavioral flexibility

Early testing showed that rapid acquisition of new targets on the standard Barnes maze was possible and can be performed with the STARR maze protocol even without adding barriers [30]. However, when challenged with a rapidly changing target, mice may switch to serial checking. The STARR maze protocol incorporates radial-arm type barriers, encouraging the mice to return to the center. By mimicking a radial arm maze and discouraging serial checking, repeat visits can be analyzed as a potential marker of working memory errors or perseveration, but without the need for food restriction [45].

We evaluated outcomes over four models with relatively similar versions of the STARR maze protocol (Fig. 8A).

For the 5XFAD experiment (Fig. 8A), the STARR maze protocol was completed more than one month following the initial spatial training, and no final probe trial was used. We found no statistical differences between the groups. This may be attributable to multiple factors, including a greater delay after the first protocol that may have decreased performance in both groups, high variability in this genotype, and fewer than the recommended number of subjects. We include the data for comparison to aid in experimental planning.

All other STARR maze experiments (Fig. 8C-E) were performed one week following initial Barnes FIELD training on the same day as the 1-week probe. In these experiments, there was a significant main effect of trial when training on the first new target, demonstrating evidence of learning (spatial + total learning). In agreement with this, aged virgin (Fig. 8L + 8P), saline-injected and DSP4-injected (Fig. 8M and 8Q) mice showed evidence of outperforming chance during the first new target trials, with unique visits lower (better) than statistical chance.

On the “reversal trials”, trial 6-9, a main effect of trial on latency was only seen in the DSP4 experiment (Fig. 8E). This experiment used our final version of the protocol, with 12 radial arms. Despite less clear evidence for rapid spatial acquisition using the learning trials, groups from all three experiments still showed statistical evidence that they favored, and therefore spatially remembered, the most recent (reversal) target.

This included control and kaolin-injected mice (Fig. 8S), aged virgin female mice (Fig. 8T), and both saline and DSP4-injected mice (Fig. 8U). We did not statistically test for preference for other prior targets, as time spent between targets is interdependent and would need a different analysis type. However, it is interesting to note for context that the post-partum group, which had high performance on Barnes Day 1-3, appeared to spend more time in the original, rather than recent target location.

### VIII: STARR maze probe for exploration vs. exploitation (win-stay vs. win-shift)

In addition to evaluation of short-term memory of the recently acquired target during the first 90 seconds of the STARR maze probe, we tested if we could identify working memory errors during full exploration of the maze. Mice were allowed to continue to explore after 90 seconds until they had visited all but one of the radial arms to measure repeat visits (errors), akin to radial arm protocols [45]. We hypothesized that after discovering the target hole was covered, a typical mouse would systematically explore the rest of the maze, making more working memory errors as more arms were explored. Indeed, we found a significant effect of “arms explored”; in both experiments, with very few errors for 4-5 arms, and increased difficulty as more of the maze is explored (Supplemental Fig. 5). Interestingly, DSP4-injected mice showed a trend towards fewer errors (Supplemental Fig. 5A), with equivalent latency to complete maze exploration (Supplemental Fig. 5B). Importantly, double checking and more time spent in the old target is considered evidence of better memory, but here would also be considered a working memory error. This highlights the potential choice between win-stay and win-shift strategies during many working memory tasks.

We therefor use the term “repeat visits”, rather than working memory errors, to highlight we do not know the motivation, although working memory errors should be more evenly distributed rather than concentrated on prior targets. We hypothesized dopamine blockade via SCH23390, a D1 dopamine antagonist, would impact working memory. Instead, we saw a trend towards reduced repeat visits (Supplemental Fig. 5E, G), even though mice were initially significantly slower to explore (Supplemental Fig. 5F, H).

Further experiments are necessary to distinguish the cause of this finding and are beyond the scope of this dataset. However, they are important to highlight here to aid in interpretation of repeat visit outcomes on radial arm or y-maze styles.

## DISCUSSION

We modified a classic, widely used, land-based spatial acquisition and retention task to better identify learning rates and improve reproducibility through use of 3D printable shuttles and a home-cage attachment that can be made by any laboratory. Importantly, our findings comparing outcome measures allow improved interpretation of the Barnes maze, even for protocols that vary from the one recommended here.

To test the utility and necessity of our 3D printed shuttles, we tested the impact of a solitary reused escape shuttle, as is the standard for many commercial systems.

Although the reused shuttle did not measurably change latency or distance outcomes during learning trials, our probe trials provide clear evidence that mice could detect the odor of the reused escape shuttle, remembering it even a week later. Thus, the presence of the odor cue during spatial learning could theoretically help or hinder learning of the spatial location in ways that might differ by experimental model. By providing evidence that mice can acquire the shuttle odor cue, we have shown that prudence is indicated when evaluating learning trials. Importantly, we showed that our 3D printed shuttles and moon caps mitigate this risk, while retaining the low-stress, naturalistic aspects of the Barnes that can make it preferable or complementary to other tasks like the Morris Water Maze.

Next, we took advantage of our random vs. spatial cue setup to compare commonly used outcome measures. For accurate statistical testing, outcome measures for each portion of the protocol should be chosen in advance of running experiments. However, as each outcome measure has distinct sensitivities and potential confounds, it is important to understand the choice. Furthermore, additional outcome measures can aid in interpretation of unexpected results.

Our study demonstrated that both latency and distance significantly improve across days, demonstrating learning. Importantly, although the spatial group did statistically outperform mice searching for a random location, based on relatively performance, much of the improvement in latency and distance across days can be attributable to other forms of learning. This is in line with other studies that have directly evaluated the Barnes maze as a spatial-learning task [46, 47]. Despite this evidence, some researchers still interpret slower latency during learning trials as specific evidence of a spatial-learning deficit, failing to consider other essential forms of learning.

Indeed, non-spatial gains may include 1) **Habituation**: Reduced novelty of the experience may initially improve latency. However, if too many trials are used, mice may shift mice towards increased exploration (unpublished observation), and habituation can differ strikingly by strain [48]. 2) **Motivational**: Running speed and perseverance may increase because of knowledge of the solution and anticipation of the reward.

Motivation may be impacted by reward pathways (e.g. dopaminergic system [49]). 3) **Procedural**: Improvements in motor efficiency through practice that help better accomplish the task. Procedural improvements, as they become automatic, are thought to be mediated by striatum [50]. 4) **Meta-learning** (learning to learn): mice learn the task structure, develop strategies, improve sensory discrimination for extra maze cues, strengthen attention and working memory, and learn to generalize [51]. Aspects of meta-learning are thought to require prefrontal cortex [52, 53].

We found that the benefit of spatial learning becomes more detectable as training duration increases. However, at the same time, many of the experimental groups show the greatest differences between groups early in training. Thus, it is essential to interpret outcomes within the context of both spatial and total learning across trials.

The finding that latency and distance outcomes are highly sensitive to more generalized learning provides an ability to screen for deficits and interpret context that should be utilized. Investigators should not restrict analysis to the probe trial or unique visits to isolate spatial learning, as they may miss non-spatial learning and motivational deficits that may drive the main differences between groups. Instead, we recommend evaluating learning trials using one holistic outcome and one spatially-restricted outcome. Specifically, based on our findings we generally recommend testing “latency to target” as the most sensitive measures of any group difference, unless there is clear concern that the mouse model has motor deficits. For spatial learning rate, we recommend “unique visits,” evaluating if they align with probe trial results to reduce risk of type 1 error. “Unique visits” during learning trials allows direct comparison with chance.

We also recommend outcomes be averaged by day in most circumstances, unless deficits are only expected on early trials. Using latency and unique visits, researchers can screen neurodegenerative models with better insight into the causes of differences between the groups.

Evaluation of several different models of dementia and aging using our modified task illustrates the utility of this careful consideration for outcome measures and use of both probe and learning trials. 5xFAD mice are a model of beta-amyloid plaque accumulation and have previously been shown to have deficits in the Y maze task [54]. Studies using different versions and protocols of Barnes maze have been used as evidence that 5xFAD mice also have deficits in spatial learning and memory. Studies have reported lower success rate to find the target, higher escape latency, working memory errors, as well as increased use of non-spatial search strategies [55–57]. While our study found evidence of slower acquisition in the 5xFAD group, it is most notable that 5xFAD mice improved to near wildtype performance with just 3 days of training, a finding that is supported by other published studies. For example, differences between 5xFAD and WT mice were no longer evident by the 2^nd^ or 3^rd^ day of training in one study, depending on outcome measure used [58]. Thus, evaluation of learning trials and intermittent probes is a compelling and useful tool to detect these clinically important early training differences. Most dramatically, total distance travelled on the probe trial showed striking differences between the groups that changed across time (Fig. 7A), suggesting motivation as a major contribution to the 5xFAD phenotype that needs to be evaluated further.

Our second model investigated outcome measures for mice that underwent intraventricular kaolin injection as a model of normal pressure hydrocephalus [26]. Prior analysis of this data demonstrated that mice with kaolin injection had increased latency to find the target during learning trials [26]. One explanation of this finding could include reduced motor speed. However, our subsequent analysis showed there were no clear differences in total distance travelled on the early training probe trials (Fig. 7B).

Furthermore, differences between the groups were also seen with distance and total visit outcomes (Fig. 5N, 5R, 5F), which are by design insensitive to motor speed.

Together, these findings suggest the differences in latency reflect differences in learning in this model, rather than motor speed. Finally, the probe trials show evidence of spatial memory at early time points and short intervals in Kaolin-injected mice (Fig. 8C). By contrast, Kaolin-injected mice do not show evidence of memory retention at 1 week, despite strong evidence in their litter-mate controls.

Our third model examined data from one-year old female mice with and without a history of pregnancy. In this case the findings highlight a difference in learning rate, with the most notable differences seen in distance travelled during the first day of learning. This is corroborated by two separate indicators: 1) evaluation of unique visits during learning trials, which shows postpartum mice differ from statistical chance on the first day, while virgin female mice perform at chance and 2) the separate 24-hour probe trial at the beginning of Day 3 shows postpartum mice have acquired the spatial location, whereas virgin mice do not show evidence of acquisition by this timepoint. Importantly, both groups show evidence of spatial learning in the 3-min short-term memory trial at the end of the third day, as well as retention at 1 week, highlighting the importance and sensitivity of examining along the learning curve to detect differences.

For our final model, we used a manipulation for which we expected no differences in spatial learning on Days 1-3, but for which we hypothesized an impact on behavioral flexibility in the STARR maze protocol. DSP4-injection is thought to specifically impact norepinephrine neurons, which play a role in multiple aspects of arousal and learning, including flexibility strategies (win-stay vs. win-shift) and tests thought to be indicative of working-memory errors [42, 59–61] Intriguingly, while in general the learning curve was similar during Days 1-3, there was evidence of faster latencies on Day 1 in the DSP4 treated group. Given the high amount of novelty present on Day 1 of training, it is possible these differences are related to differential response to novelty rather than enhanced spatial learning perse, and further investigation of this is ongoing.

While place-preference near the learned target can test retention, the probe trials can also provide important information about motivation and flexibility. We found that total distance travelled on the probe trial can be an important measure to help provide context to our other results. <u>Total</u> distance travelled on the probe is differentially influenced by multiple factors: hyperactivity/hypoactivity, apathy/perseverance, exploration/fear, as well as the declarative memory of the task. Across our model experiments, there were differences in distance travelled on the probe trials, and these differences tended to change gradually across training. It was notable that these differences were not evident in standard open field testing (when available). For example, kaolin-injected mice showed the same distance travelled in open field [26], but striking differences during the probe trials for one-week retention, suggesting motivation, rather than motor impairment as a likely cause. Notably, saline-injected mice increased their distance travelled across training, while DSP4-injected mice did not show this effect, also leading to a group difference during the one-week retention trial.

It is important to consider how this context impacts interpretation of spatial learning percentages on probe trials. For instance, consider a hypothetical mouse that goes to the target, stops, and remains there for the entire duration. This mouse would be labeled as having “strong” spatial memory using percentage of time spent but is arguably abnormal in behavior; they would fail to explore alternative hiding places in the face of a blocked tunnel. Similarly, differences in activity can alter percentage time at sector and may or may not relate to spatial memory. For example, male and female mice performed similarly for latency and distance (not shown) but showed a statistical difference from one another during the 3-min short-term memory trial (Fig. 4D), even though both perform better than chance. It is for this reason that we recommend evaluating probe trials with a “yes or no” statistical test against chance for the entire group, along with visual inspection of the data. It is essential to understand that differences in the magnitude above chance likely reflect factors other than spatial memory.

Finally, we developed a test for rapid acquisition and behavioral flexibility that can be added after any traditional Barnes protocol. As noted above, group differences are often detectable in early training but become less apparent with extended training or overtraining. We designed and tested the one-day Spatial Training and Rapid Reversal: STARR maze as an add-on to our Barnes FIELD protocol. It allows assessment of spatial location acquisition over a rapid time frame, using four trials at a location distinct from the one originally acquired, followed by four trials at a third location. We have previously published evaluation of a Parkinson’s model of cortical synuclein using this task without barriers, showing slowed learning of the reversal target in this model [30]. In that study, we were able to repeat the protocol multiple times, including in the setting of injection with lipopolysaccharide (LPS). Here, we tested the use of barriers to prevent serial checking and evaluate exploration efficiency and repeat entries, akin to a radial arm maze.

The STARR maze behavior flexibility protocol adds several important new features. Following the one-week probe (trial 1), barriers are introduced which add novelty to the environment and disrupt previously acquired search strategies. The first target change (trial 2-5) challenges mice to rapidly acquire a new location in the setting of this novelty. The second target change (trial 6-9) more purely tests rapid acquisition, as the novelty has started to fade. The shift from outperforming chance on trial 5, to meeting chance on trial 6 (for unique visits) provides evidence that some groups learn the task and then have more difficulty with the new target (Fig. 8J, 8K, 8L, 8M). Interestingly, some mice that had performed well on Days 1-3 (e.g. aged postpartum females) showed a tendency towards favoring the original target when evaluated on the probe, suggesting the original target location may be retained in these mice.

Most excitingly, DSP4-injected mice, which had generally performed similarly on Days 1-3 of the task compared with saline-injected mice, showed more clear changes with this behavioral flexibility protocol, a major goal of our behavioral optimization. These mice were slower to find the new target on average during trial 2-4, and while they did outperform chance on average, evaluation by trial suggests they required more trials compared with saline-injected mice. Comparing the probe trials suggests a similar finding; the DSP4-injected mice acquire the most recent target, but there is evidence that precision is reduced. It is important to highlight that although the third target switch was clearly more difficult (few groups across experiments outperformed chance for unique visits during acquisition), most groups favored the more recent target during the probe trial. This provides evidence that wildtype mice that have received this training protocol can rapidly acquire a new spatial location. Therefore, the key to interpreting outcomes is examining them together in context for evidence of converging results.

Finally, evaluation of exploration on the final probe provides the opportunity to test for repeated visits. Although we initially designed the extended probe trial of the STARR maze to identify “working memory errors”, it is also possible that repeat entries represent a win-stay vs. win-shift pattern, rather than an error. This is highlighted by our DSP4 experiment; over the 90-second probe, control mice spent more time in the specific target arm than other arms (Fig. 8U, 8Y), akin to “win-stay”. When the mice were allowed to continue exploring, this may also cause a tendency for control mice to be slower to explore the entire maze and making significantly more “errors” (Sup Fig. 3). This type of context is essential for understanding impacts on executive function. For example, the locus coeruleus is implicated in win-shift vs. win-stay strategy, but has variably been implicated in working memory [60, 62–65]. Further work is needed to determine why DSP4 causes the outcome seen here. Regardless, it is important to note that without the context of both phases of the protocol (Barnes FIELD and STARR maze), one might incompletely conclude DSP4-injected mice were specifically better at spatial memory acquisition.

### Limitations

Limitations and remaining gaps are addressed throughout the manuscript. Of note, protocols were adapted and modified over the course of multiple years, with each cohort run under the hosting laboratory’s supervision by separate investigators adapting to the needs of their experiments. Thus, caution must be taken comparing between cognitive models presented, especially since models were not interleaved and control groups show striking differences. Similarly, we cannot rigorously test all possible permutations for the protocol when creating final recommendations, and limitations to how these choices were made are noted in the detailed protocol.

Human-run mouse behavioral studies show generally higher variability compared with repeated, automated testing. However, operant chambers and in-cage testing are costly, achieve reduced variability with overtraining, and often require reward or punishment-based strategies. A major problem in behavioral literature in general is publication bias; researchers leave out variable behavioral data because reviewers expect to see tight outcome measures among groups that are unrealistic for animal behavior. This self-fulfilling prophesy leads to more publication bias. Thus, while variability in our outcomes is a limitation, and highlights when larger sample sizes are necessary, our representative data also represents a key benchmark important to the field. Although human-run screening studies are usually more variable, we believe there will always remain a benefit for naturalistic tasks observed by humans to allow direct discovery of unexpected behavioral findings. As with direct interaction with human patients, there remains an important role for direct observation in mouse models.

A second limitation to our study was that we only ran the final version of protocol on the final model (DSP4) included in the manuscript, while earlier models used 10-arm versions or lacked specific probe trials. The STARR maze protocol required years of optimization to get to the current protocol, and many of the choices balanced major trade-offs. For example, one version of the maze required mice to push full caps off each hole, reducing serial checking and thus adding statistical power. Unfortunately, while control mice learned this variation in about two days, impaired mice often failed to remember how to move the caps by the next day, preventing evaluation of any of the other variables. Similarly, an early iteration of the STARR maze protocol had rapid reversal learning on Day 4 (no barriers) and added barriers to identify repeated entries on Day 5. Although the protocol was successful and less variable, very few labs were able to run a 5-day protocol with the space and time allotted. For this reason, we attempted to optimize a combined 4-day add-on protocol using both reversal and barriers. This add-on protocol may be further optimized in future research as we evaluate additional models of cognitive impairment.

Refining rodent behavioral paradigms remains crucial for both basic and translational research. As drugs or interventions need to prove efficacy in animal models before they can be considered for human use [66], it is imperative that outcome measures are well understood.

Despite the mentioned limitations, we present evidence for essential modifications to the classic Barnes maze to improve and clarify interpretation of outcome measures. Our findings have important implications for interpretation of both old and new versions of spatial acquisition tasks and provide steps to build this accessible cognitive screen at low cost. These features will improve rigor and reproducibility across neuroscience laboratories using behavioral outcomes.

