## Supplemental Methods for "Renovating the Barnes maze for mouse models of Dementia with STARR FIELD: A 4-day protocol for learning rate, retention, and cognitive flexibility"

**Barnes FIELD Procedure: Step-by-step**

1) Don lab coat and gloves. Make all needed cleaning solutions. Turn on flood lights. Preclean maze. Fully set up all shuttles with one shuttle below each hole, and all holes blocked by grates except the target hole. A moon-shaped cap should block each hole from visual view of the center. Turn on video recording and test maze rotation alignment. Set up naming scheme for files. Write name of **mouse**, **trial number**, and **target** on a white board in field of view of camera. For the complete protocol, there will be four targets: Habituation, Barnes(Day1-3), New, Reverse. Each target should be selected to be at the center of a quadrant, and two groups should be counterbalanced with opposing targets.
2) **Habituation**: For the habituation trial, bedding from the home cage is placed into the open habituation target and near the stairs. Following the habituation trial, this shuttle is removed and completely washed and replaced with a new trial. Bedding is only used on the habituation trial.

**Learning Trials**: Following one habituation trial, the spatial target is changed to a new quadrant, and remains in place for all subsequent trials on Days 1-3. Although the maze platform turns to reduce odor cues, a single spatial location contains an open shuttle.

**Probe Trials:** For probe trials, all clean shuttles are in place and blocked by grates. The maze is turned as with a standard learning trial. Probe trials last 90 seconds.

3) For each trial the mouse should start in its original home cage (not a clean cage). Cage-mates should be placed in a separate clean cage away from the task until testing is complete.
4) Place the mouse into the funnel, cover with the transport platform, then gentle invert the funnel so the mouse is standing on the platform.
5) Place the platform, mouse and funnel in the center of the maze. Gently slide out the platform taking care not to place the funnel on top of the mouses tail. Leave the funnel in place.
6) Start the video recording and wait 10 seconds. Then, lift the funnel and step out of the testing area promptly. Monitor the mouse by video and wait until the mouse enters the target hole. Always remain quietly in the same position during the trial.
7) Only for probe trial: wait the listed time, adding 10 seconds if using the video recording clock and then remove the mouse. Use the funnel/platform in reverse to remove the mouse. Use caution during the probe trials not to handle the mouse roughly as this may increase variability, causing increased fear-related learning in some mice. Trained mice can be scooped by hand, less well-handled mice can be picked up carefully by the base of the tail. Take care to use the same method for all mice in the experiment.
8) Stop the video recording.
9) For learning trials: Once the mouse enters the target, cover the target hole with a cap to prevent the mouse from leaving its shuttle. Bring the home cage into the testing area. Attach the shuttle to the home-cage via the docking tunnel and cover the home-cage with a drape to further incentivize the mouse’s return by making the cage dark. Remove the shuttle door to allow the mouse entrance to the home-cage.
10) While the mouse is getting “rewarded” (either sitting in the shuttle or making its way down to the home cage), wipe the maze with 70% ethanol solution. Once you’re done cleaning the maze and the mouse is in the home cage, disconnect the docking-tunnel from the shuttle and home-cage, and remove the shuttle from the maze. Roll the cart/home-cage out of the training area and back to its idle position. Clean the shuttle per protocol and put it to dry. Choose an unused, clean, dry shuttle to put back under the maze.
11) Rotate the maze (3 – 4 holes) clockwise, taking care not to use a position that has been used during this training session. Example platform positions can be found in Figure 2. Remove the grate from the shuttle that is now at Spatial Location A (for this example*). Make sure all other shuttles are blocked and fully pushed into position. This takes about 2-3 minutes (inter-trial interval). Using the starting funnel, place the mouse in the center of the maze, covered by the funnel. Repeat steps 3-9 above for each trial.
12) For subsequent mice: Wash the maze with an appropriate agent. Counterbalance your target escape zone across experimental groups (2 versions should be sufficient, greater numbers will lead to more human error).

**STARR maze Procedure: Step-by-step**

1. Run either the one-week probe or a habituation trial as trial 1 (no barriers).
2. Add the barriers, using the barrier holders.
3. Follow steps 1-9 of the Barnes FIELD procedure with the following notable changes:
4. For trials 2-5, the spatial target (open shuttle) is located in a new quadrant (NEW).
5. The maze is turned 3-4 holes clockwise during trials 2-4). After trial 5, do not turn the maze (the location of the target (open) shuttle will change instead).
6. For trials 6-9, the spatial target is set to a new quadrant (REV). After each of trials 6-9, turn the maze counterclockwise 3-4 holes, taking care not to use a previously used position. Position examples for a 12 hole maze can be found in figure 2.
7. For trial 10: Run a probe trial with the barriers all in place. IMPORTANT: Stop the trial when the mouse visits **at least n-1 arms** and **at least 10 minutes have elapsed**, where n is the number of holes on the maze. This gives both an exploration distance over a set interval, and a statistical estimate of number of working memory errors over a set amount of explored maze. Mice do not usually begin to make working memory errors until they have visited around 5 arms.

Each target chosen for the STARR maze should have two adjacent holes/arms beside them that have not been used for target acquisition (Fig. 2). When choosing targets and counter-balancing, ensure adjacent slices will not overlap with previous targets/or target quadrants. This is simplest in a 12-hole maze, where each target can be at the center of one quadrant.

**Additional Features:**

**Aversive Sound:** Some laboratories recommend a loud sound in addition to bright lights to drive mice to enter the hiding space, which is turned off when the mice enter. Due to risk of increased experimenter error (timing of turning off cue when hole entered) as well as experimenter fatigue, annoyance, and interference in other nearby experiments, we have not tested this option. Instead, we use the home cage as a motivational factor instead of an aversive factor.

**Intramaze or alternative cues**: Early pilot experiments showed mice could be trained to a single cue as a control (a 3D-printed structure used for novel object training placed by the hole). By contrast, use of a different object at each hole was difficult and required longer training periods. Training to shifting cues increased difficulty and variability, as some mice have more difficulty switching cues. These protocols have not been sufficiently optimized to recommend.

It is always important to remember that multiple extra-maze cues, in addition to visual, are available to the mouse. These include light gradients, scent gradients, heat gradients, vibration and sound gradients. Many spaces may contain noises that are inaudible to humans that may be also used as spatial cues or even be aversive. These cues create a spatial location, making this task distinct from some protocols where only visual cues are used and moved during each trial.

**Maximum training days**: Pilot studies using daily STARR maze training, with new spatial locations each day suggest that mice can be trained/tested for > 7 days in a row with good performance. However, after about 7 days, mice may become too habituated and spend more time exploring the surface of the maze rather than prioritizing finding the target. By contrast, we have used intermittent evaluation (1-3 days at a time, months apart) in prior published studies [1].

**Troubleshooting:**

1) If the mouse falls or jumps off the apparatus during testing, **keep recording**; promptly enter the testing area and return the mouse to the center of the apparatus.
2) If the mouse is resistant to leaving the shuttle, remove the tunnel and place the entire shuttle (mouse still inside), inside the home cage. Usually, the mouse will then leave on their own or can be gently coaxed out.
3) If the mouse is resistant to leave the docking-tunnel. Place the entire tunnel in the cage. If they do not then leave on their own, the tunnel is designed with openings to allow the mouse to be coaxed out gently.
4) Shuttle drying: Printing ~30 shuttles allows the shuttles time to dry between mice, allowing continuous running. It is recommended that you print additional copies of all items (~2-3 escape tunnels), >30 copies of all other items (caps, grates, shuttle doors).
5) Mice cannot find the shuttle: Typically used protocols recommend stopping the trial after 3 minutes [2] if the mouse fails to find the target. Unfortunately, ending the gfidl due to a set cutoff time prevents use of “distance”, “visits”, or “unique visits” as primary outcome measures because a mouse that doesn’t move much will have a “short” (better) distance. For these reasons, we recommend allowing mice unlimited time to locate the target. However, we have found some mice (with specific treatments or conditions) fall asleep on the maze when they fail to find the target. In these cases, we have used experimenter encouragement to allow the mice to complete the maze. This is not ideal (experimenter involvement increases variability and potential bias), but we found on balance provides slow or anxious mice the opportunity to learn the spatial target. We have developed a printable intervention protocol that can be used with difficult mouse models (supplemental Fig 3). It is important statistically that the investigator does not encourage the mouse to the target hole, only to the center. Failure to find the target hole is a rare occurrence in most mice, but may be expected for some severe phenotypes or occasionally when running a large cohort of mice. As with all studies, the experimenter must use some judgement in adapting the protocol to severe phenotypes to avoid misattributing motivational, exploration or mood abnormalities to cognitive (memory, learning, attention, etc.) deficits.
