## Supplemental Figures 1-6 for "Renovating the Barnes maze for mouse models of Dementia with STARR FIELD: A 4-day protocol for learning rate, retention, and cognitive flexibility"

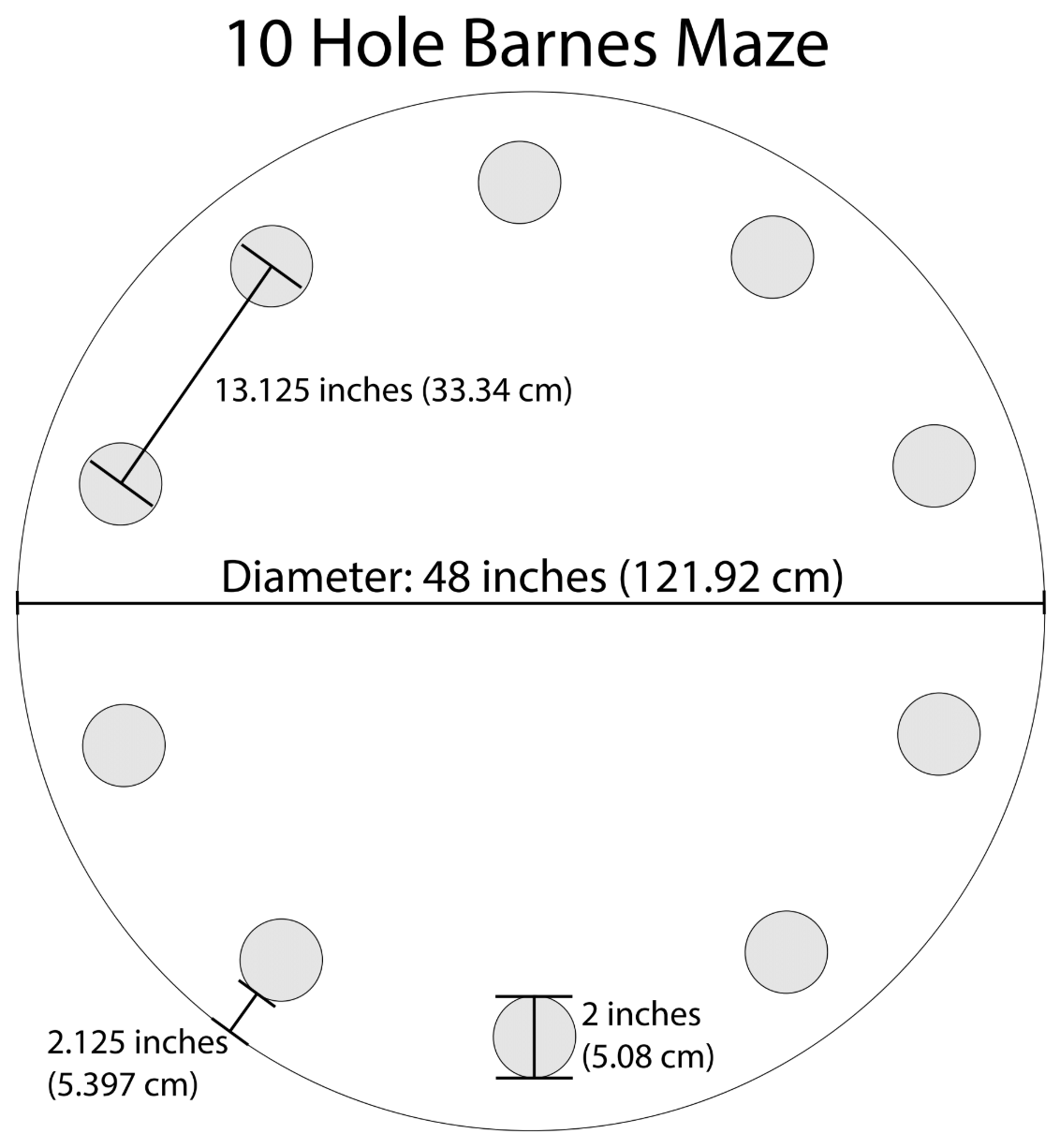

**Supplemental Figure 1: Schematics for a 10-hole Barnes maze.** Schematic for production by plastic manufacturer. The use of 10-holes is an example, as it can include as many or as few holes as the experiment requires. This diagram can be used request custom cutting for a high density polyethylene (HDPE) platform, typically around 0.635 cm thick.

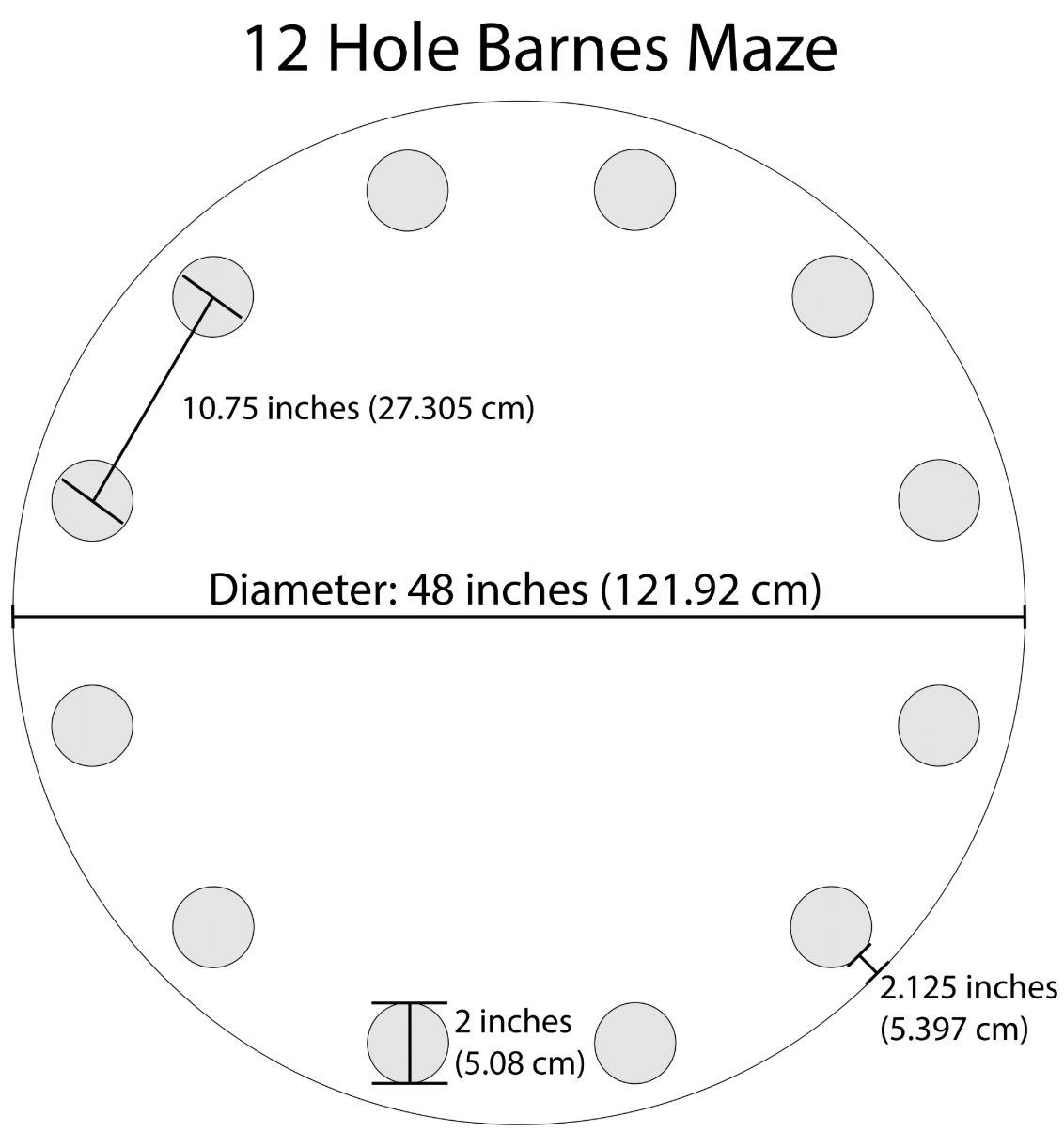

**Supplemental Figure 2: Schematics for a 12-hole Barnes maze.** 12-hole Barnes maze schematic for production by plastic manufacturer. 12-holes is the recommended number used in our final protocol. This diagram can be used request custom cutting for a high density polyethylene (HDPE) platform, typically around 0.635 cm thick.

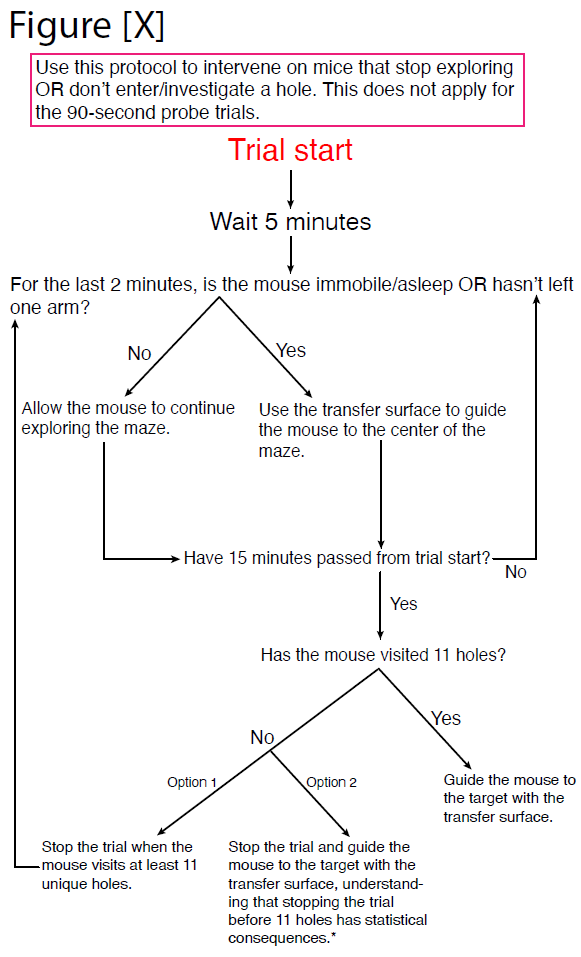

**Supplemental Figure 3: Printable intervention protocol.** Most mice do not require intervention. However, some severe models may impact arousal and cause mice to fall asleep or stop exploring. Investigators can choose to end trials after 15 minutes or allow the mouse unlimited time to explore “h-1” unique holes of the maze (where h is the number of holes). The last probe on Day 4 (trial 10), should be > 10 minutes long AND allow the mouse to visit enough holes. Waiting until the mouse has visited at least “h-1” holes is statistically preferred when comparing between groups, unless time becomes excessive.

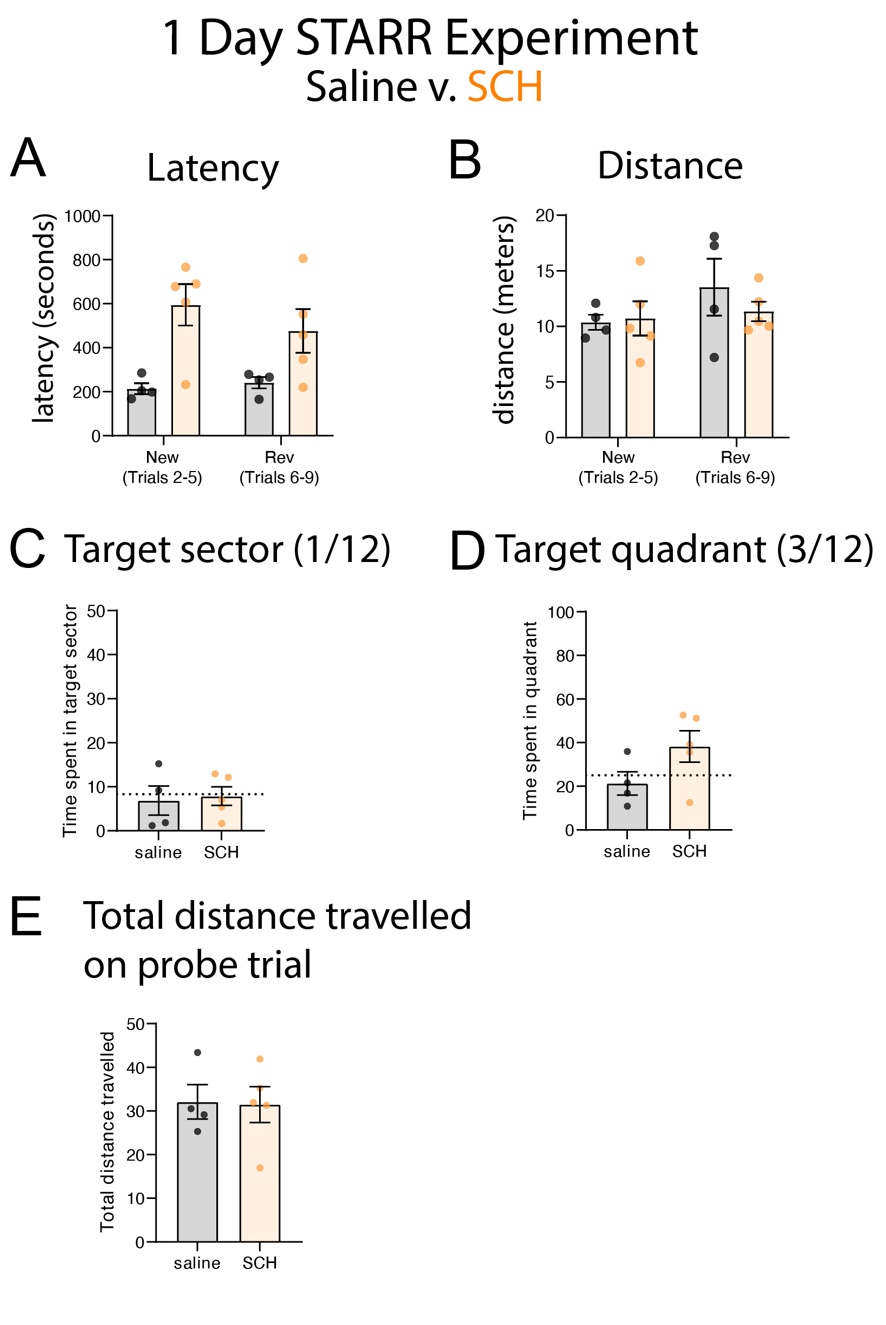

**Supplemental Figure 4: Latency and distance outcomes diverge under certain circumstances.** While latency (A) and distance (B) outcomes usually concur, manipulations with motor impact or that cause immobile spells (freezing or sleep) may only be revealed with latency. In a single-day STARR maze, we compared saline injection vs. SCH23390 (a dopamine receptor antagonist). SCH23390 increased latency during STARR maze trials, without impacting distance to target. Although this experiment helped demonstrate how latency and distance can differ as outcome measures, we also found performing a single-day STARR maze (without the preceding 3-days of normal spatial training) does not provide sufficient time for mice to “learn to learn”. Specifically, saline mice did not spend increased time in the most recent target sector (C) or quadrant (D). However, this experiment was also underpowered for this outcome (n = 4, 5). For this reason, we recommend the STARR maze be performed after the standard Barnes maze.

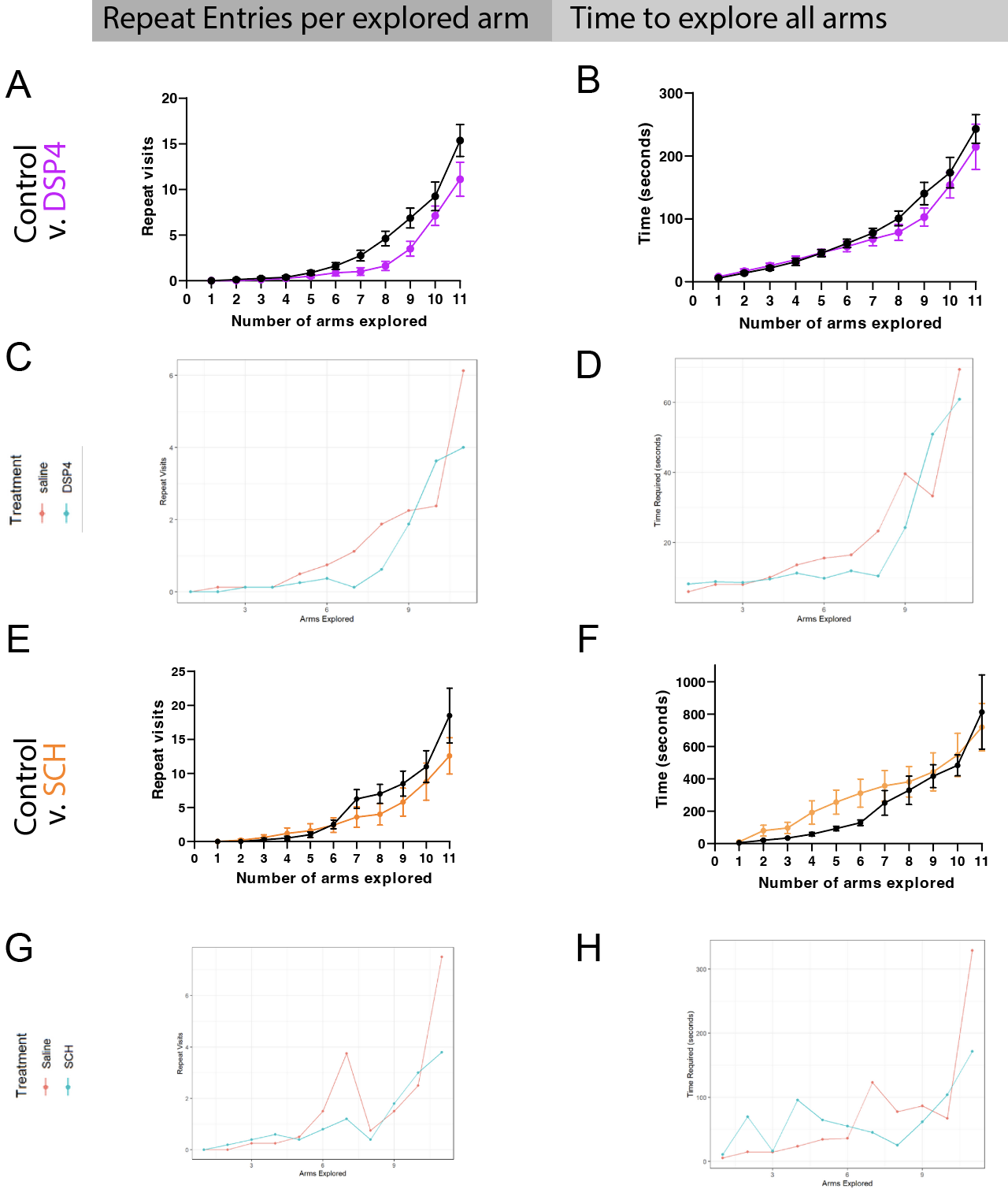

Supplemental Figure 5: **Repeat Visits**: Number of repeat visits as a function of total arms explored is a measure of exploration efficiency. While “repeat visits” are usually considered “working memory errors”, they may also represent a “win-stay” exploratory strategy, especially in the context of prior training. Repeat visits were defined as returning to the center and visiting another arm, before returning to a previously visited arm. Repeat visits were graphed cumulatively (A) but analyzed using number of new errors for each new arm explored (C). There was a main effect of arms explored (p< 0.001), demonstrating mice started making more errors as more arms were explored. There was a trend towards fewer repeat visits in the DSP4 treated mice (Fig. A, p = 0.079, interaction: p= 0.182, GLMM with Negative binomial). For SCH treated mice, there was no difference in number of errors (main effect arms: p < 0.001, SCH: p =0.833. interaction: p = 0.516). **Exploration efficiency**: The cumulative (B, F) and net (D, H) time required to explore each new arm of the maze is also a measure of exploration efficiency. No difference between groups was seen for DSP4 treated mice (Fig. B, GLMM with log-transformed Gamma model, main effect arms: p<0.001, DSP4 p = 0.569, int: p = 0.223). By contrast, mice treated with SCH were significantly slower to start exploring arms of the maze (Fig. F-H, main effect arms: p < 0.001, SCH: p = 0.002, interaction: p<0.001).
